# Isolation of Biologically Active Extracellular Vesicles-Associated AAVs for Gene Delivery to the Brain by Size Exclusion Chromatography

**DOI:** 10.1101/2023.05.30.542901

**Authors:** Carina Henriques, Miguel M. Lopes, Patrícia Albuquerque, David Rufino-Ramos, Laetitia S. Gaspar, Diana Lobo, Kevin Leandro, Ana Carolina Silva, Rafael Baganha, Sónia Duarte, Casey A. Maguire, Magda Santana, Luís Pereira de Almeida, Rui Jorge Nobre

## Abstract

Extracellular vesicles-associated adeno-associated viral vectors (EV-AAVs) emerged as a new opportunity for non-invasive gene therapy targeting the central nervous system (CNS). However, in previous reports, only AAV serotypes with known ability to cross the blood-brain barrier (BBB) have been used for EV-AAV production and testing through non-invasive strategies. In this work, we aimed at optimizing a size exclusion chromatography (SEC) protocol for the production and isolation of natural and biologically active brain-targeting EV-AAVs, that could be applied to any AAV serotype and further used for non-invasive gene delivery to the CNS. We performed a comparison between SEC and differential ultracentrifugation (UC) isolation protocols in terms of yield, contaminants, and transgene expression efficiency. We found that SEC allows a higher recovery of EV-AAVs, free of cell contaminating proteins and with less *solo* AAVs than UC. Remarkably, SEC-purified EV-AAVs also showed to be more potent at transgene expression than *solo* AAVs in neuronal cell lines. EV-AAVs exhibited the ability to cross the BBB in neonatal mice upon intravenous administration. In conclusion, SEC-purified brain-targeting EV-AAVs show to be a promising gene delivery vector for therapy of brain disorders.

**Graphical Abstract:** 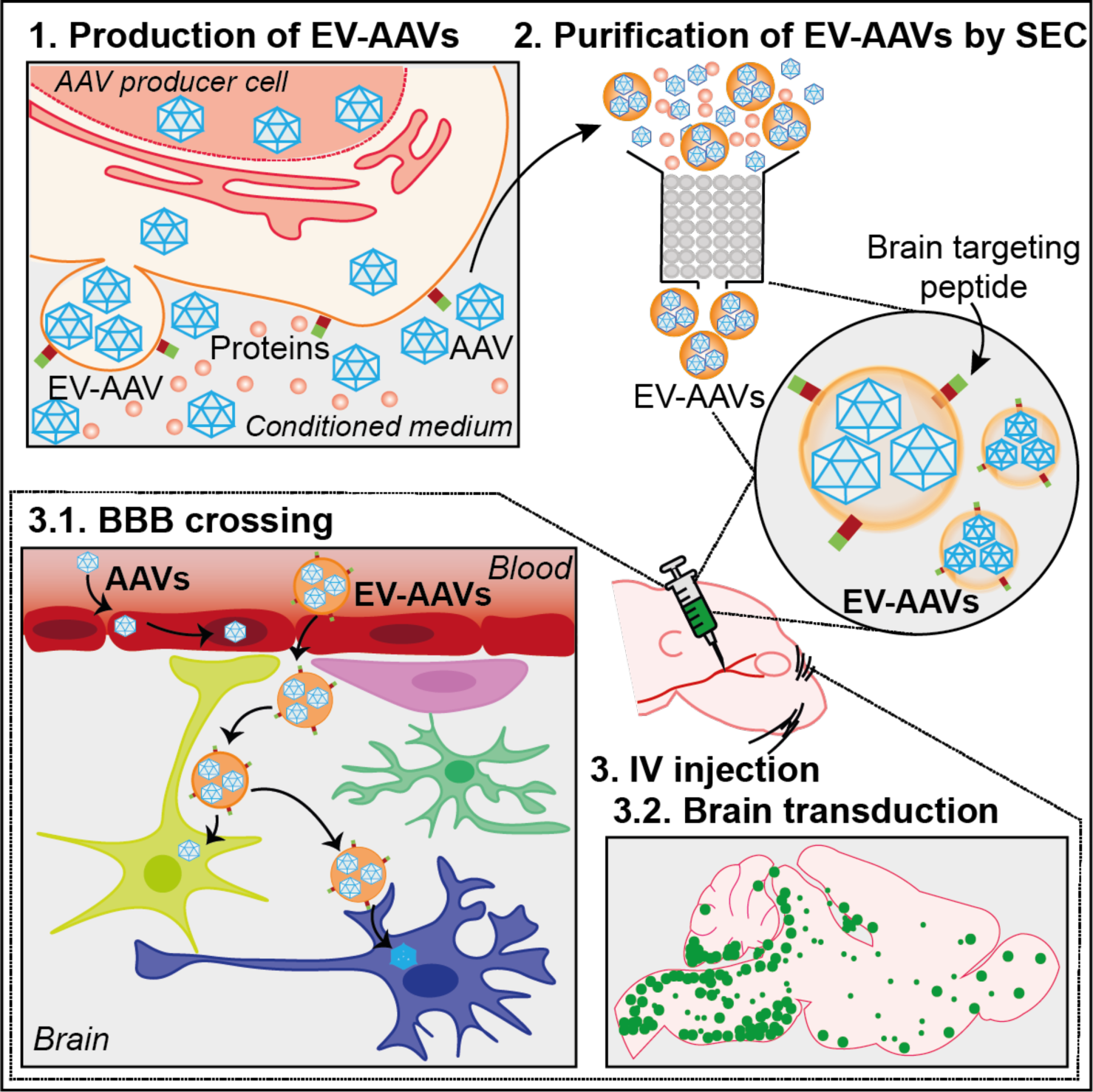

During the production of AAV vectors, a small percentage of AAVs is secreted in association with extracellular vesicles, named “EV-AAVs”. EV-AAVs can be efficiently isolated by size exclusion chromatography (SEC). When intravenously injected in mice, brain targeting EV-AAVs can cross the blood brain barrier (BBB) and transduce neuronal cells.

## 1. Introduction

In the last few years, adeno-associated viral vectors (AAVs) have been the vector of choice for the delivery of gene products to the human body, specially to the brain^1^. Wild-type AAVs are small viruses, with a big repertory of serotypes with different tissue tropisms^2, 3^. Currently, six AAV-based products have been approved for the treatment of human diseases in Europe and the USA^4–9^. However, new difficulties have emerged with clinical translation and use of these vectors^10^. A major challenge is the high prevalence of pre-existing neutralizing antibodies against AAV capsids in humans, related to previous AAV infection or cross-reactivity between different AAV serotypes^11^. In addition, for central nervous system (CNS) disorders, high doses of systemically delivered AAVs need to be used to reach the brain, even when AAV serotypes known to cross the blood brain barrier (BBB) are used (e.g. AAV8 and AAV9). Moreover, upon intravenous (IV) administration, most AAV serotypes show tropism to peripheral tissues (e.g. liver) causing the risk of undesired expression in off-target tissues and toxicity^12^. To circumvent these limitations, a significant effort has been made to develop new AAV capsids able to avoid neutralization, efficiently cross the BBB and transduce specific regions or cell populations of the CNS, while simultaneously maintaining a low transduction of peripheral organs^10, 13^.

During the production of AAVs, a portion of these vectors were found to be secreted into the media in association with extracellular vesicles (EVs), the so-called EV-AAVs^13^. Several reports have demonstrated that EV-AAVs are not only able to achieve a higher cell transduction than AAVs^14–19^, but also to circumvent antibody neutralization^13–16, 19^ and biological barriers such as the BBB^14, 20^. However, only AAV serotypes with already known ability to cross the BBB upon IV injection were used for the development of EV-AAVs targeting the CNS. It remains unclear whether AAV serotypes without natural capacity to cross the BBB can reach the brain in association with EVs. Moreover, there is still the need for an isolation protocol that allows the recovery of pure and biologically formed EV-AAVs with potential for scale-up and clinical translation.

The present work aimed to optimize and validate a protocol for the isolation of purer and biologically active EV-AAVs, able to cross the BBB and transduce the CNS, with the potential to be translated into clinical applications (Graphical Abstract).

## 2. Results

### 2.1. Size exclusion chromatography allows the isolation of pure and biologically active EV-AAVs

Size exclusion chromatography (SEC) has been shown to be a simple and fast strategy to isolate highly pure, intact and functional EVs, with potential application in clinical setting^21–23^. In the present work, we aimed at optimizing a SEC protocol for the isolation of EV-AAVs, based on a protocol previously developed by our group to isolate EVs from plasma^24^.To achieve that, we took advantage of mosaic AAV vectors carrying capsid proteins from AAV1 and AAV2, thus combining the properties of both AAV serotypes that, despite their neurotropic features, are not able to efficiently cross the BBB^25^. In our protocol, HEK293T cells were transfected with the four plasmids for AAV1/2 production: (I) a plasmid encoding the enhanced green fluorescent reporter gene (GFP), under the action of chicken beta-actin (CBA) promoter placed between the ITRs sequences (pITR), (II, III) two plasmids containing the wild-type (WT) AAV genome rep and serotype 1 and 2 cap ORFs and (IV) a helper plasmid^26^. A fifth plasmid encoding for Rabies virus glycoprotein peptide (RVg) was also used to target EV-AAVs to the brain^14^. The conditioned medium from AAV-producer cells was then collected at 48 h and 72 h post-transfection (Figure 1.A), pre-cleared by low-speed centrifugations and concentrated using ultrafiltration (UF) with a molecular weight cut-off (MWCO) of 10 000 kDa to reduce EVs loss^27^ (Figure 1.B). The filtered concentrated medium was then applied into an agarose-based SEC column and 30 fractions (F) of 0.5 mL were collected individually for posterior analysis (Figure 1.C).

**Figure 1-.**
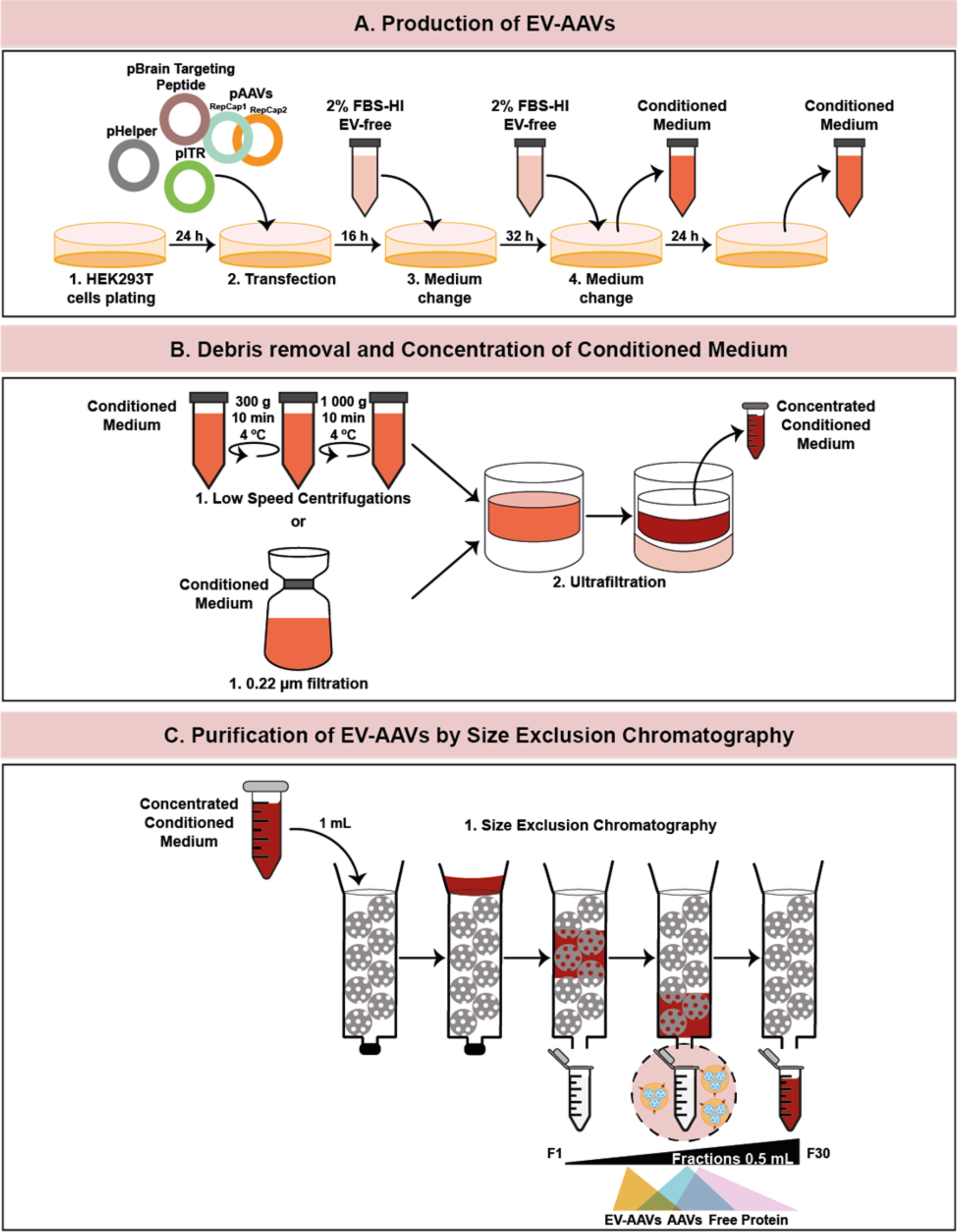
Workflow summarizing the protocol for the production and isolation of brain targeting EV-AAVs by SEC

To identify the SEC fractions where most proteins eluted, we performed a western blot on the 30 collected fractions and assessed total protein by staining membranes with Ponceau S solution. Proteins were mainly detected between F12 and F26, with a peak at F18 (Supplementary Figure 1.A and Supplementary Figure 1.B). The same profile was obtained when proteins were quantified using the bicinchoninic acid assay (BCA) (Supplementary Figure 1.C).

To find out in which SEC fraction(s) EVs were eluted, we used the marker flotillin 1 (FLOT-1). This marker is expressed in both small and medium/large EVs allowing us to identify the range of fractions containing both entities. As shown in Figure 2.A and Figure 2.B, FLOT-1 was firstly detected on F7, and showed a peak between F8 and F11. Calnexin (CNX), a typical cellular marker, was not detected between F7 to F13, indicating the absence of cell contamination. Nanoparticle analysis by NTA confirmed the presence of EVs sized from 70 nm to 130 nm in F7-13. In these fractions, we also found a gradual decrease in size mode and in the nanoparticle/protein ratio (Figure 2.C-E). Negative stain Transmission Electron Microscopy (TEM) also confirmed the presence of intact EVs between F7 and F12 (Supplementary Figure 3).

**Figure 2-.**
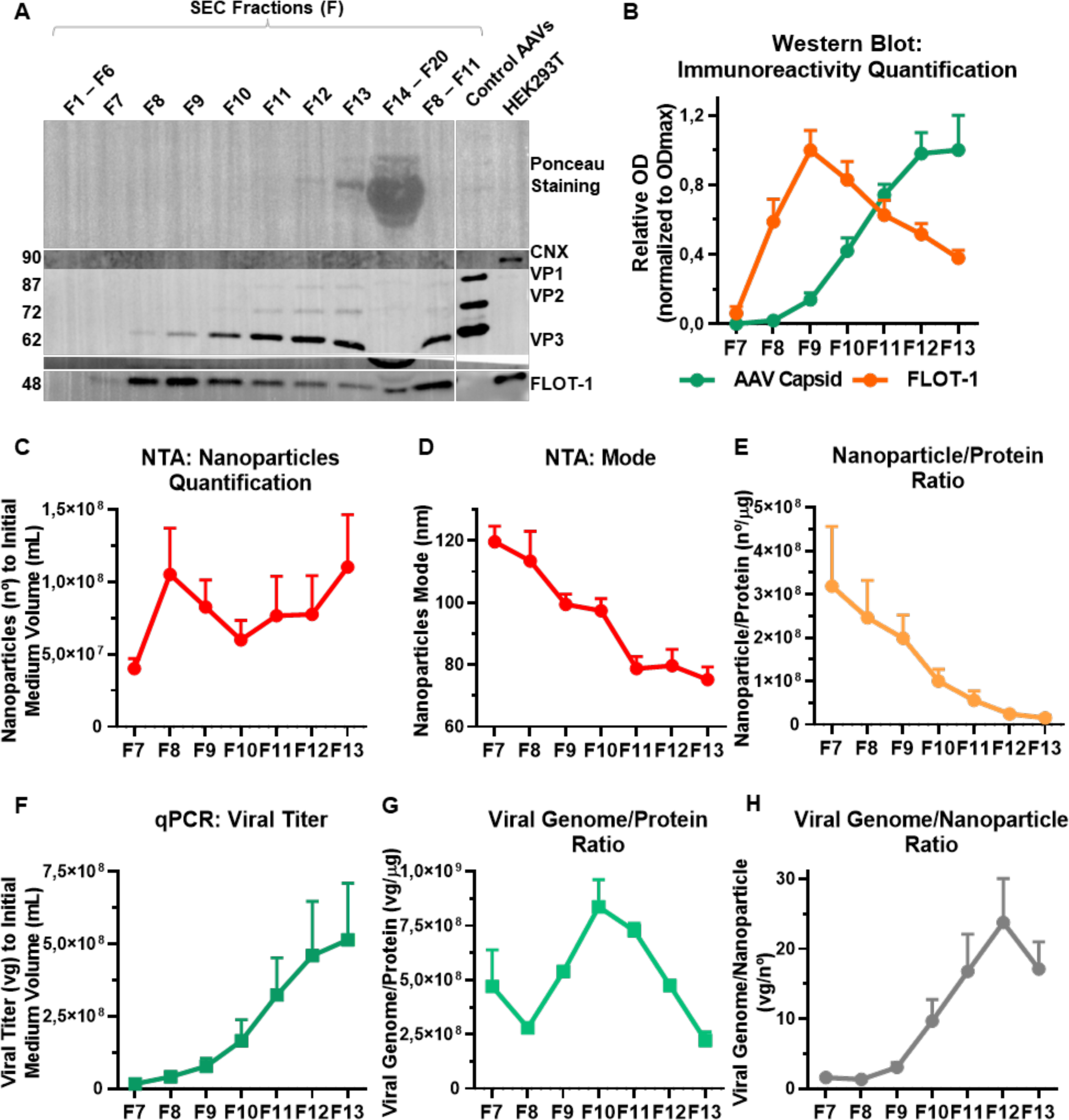
EV-AAVs can be separated by SEC

The AAV elution profile was also evaluated by western blot. AAV capsid proteins (VP1, VP2, and VP3, that assemble in a 1:1:10 ratio) were firstly detected on F8 and showed a gradual increase in F9-F13 (Figure 2.A-B and Supplementary Figure 2). Viral genome quantification by qPCR also showed a gradual increase in viral titer along these fractions (Figure 2.F). While the viral genome/protein ratio oscillated between F9-13, the ratio between viral genomes and nanoparticles (viral genome/nanoparticle) gradually increased between F7 and F12 (Figure 2.G-H).

To test the transgene expression efficiency of the most promising SEC fractions regarding the presence of EV-AAVs, Neuro 2A (N2a) cells were independently transduced with F7 to F13 at a MOI of 10 000 viral genomes/cell (Figure 3.A). Transduction was qualitatively evaluated through the analysis of GFP expression by immunocytochemistry (ICC) and quantitatively by Fluorescence-Activated Cell Sorting (FACS). F7 to F11 showed to be the most potent fractions compared to control cells, with F8 being the most potent (23.08% ± 1.69% GFP-positive cells) (Figure 3.B-D and Supplementary Figure 4).

**Figure 3-.**
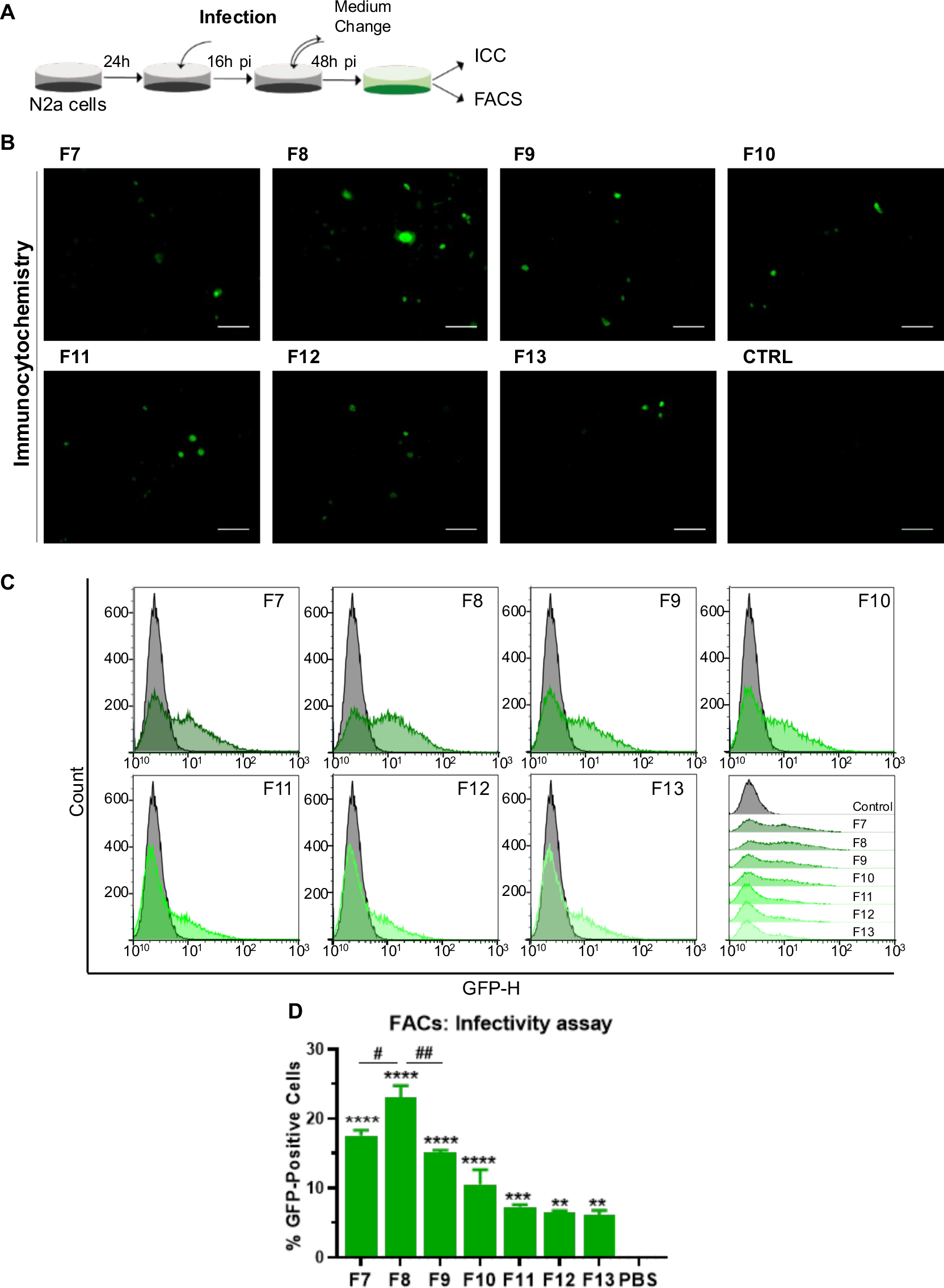
EV-AAVs (F7 to F11) are the most potent fractions in neuronal cells

Taking these results into account, we infer that the purest and biologically active population of EV-AAV1/2 coated with RVg (referred in our work as EV-AAV) is collected on SEC fractions 7 to 11. However, because of the low viral titer and EV amount observed in fraction F7 (Figure 2.A-C and Figure 2-F), we chose to exclude this fraction and proceed only with fractions F8 to F11 in the following experiments.

### 2.2. SEC allows a better purification of EV-AAVs compared to UC

Once the SEC protocol was optimized, we performed a side-by-side comparison of SEC and the standard protocol of differential centrifugation (UC)^28^. Cell medium was collected at 48h and 72h post-transfection of HEK293T cells with the same five plasmids for AAV1/2 production and RVg expression mentioned above. Cell debris were removed by low-speed centrifugations. Half of the medium was then purified by UC (UC-100k) and the other half by our SEC protocol (Figure 4.A). EV-AAVs isolated from the medium by UC and SEC were then characterized using microBCA, NTA and qPCR.

**Figure 4-.**
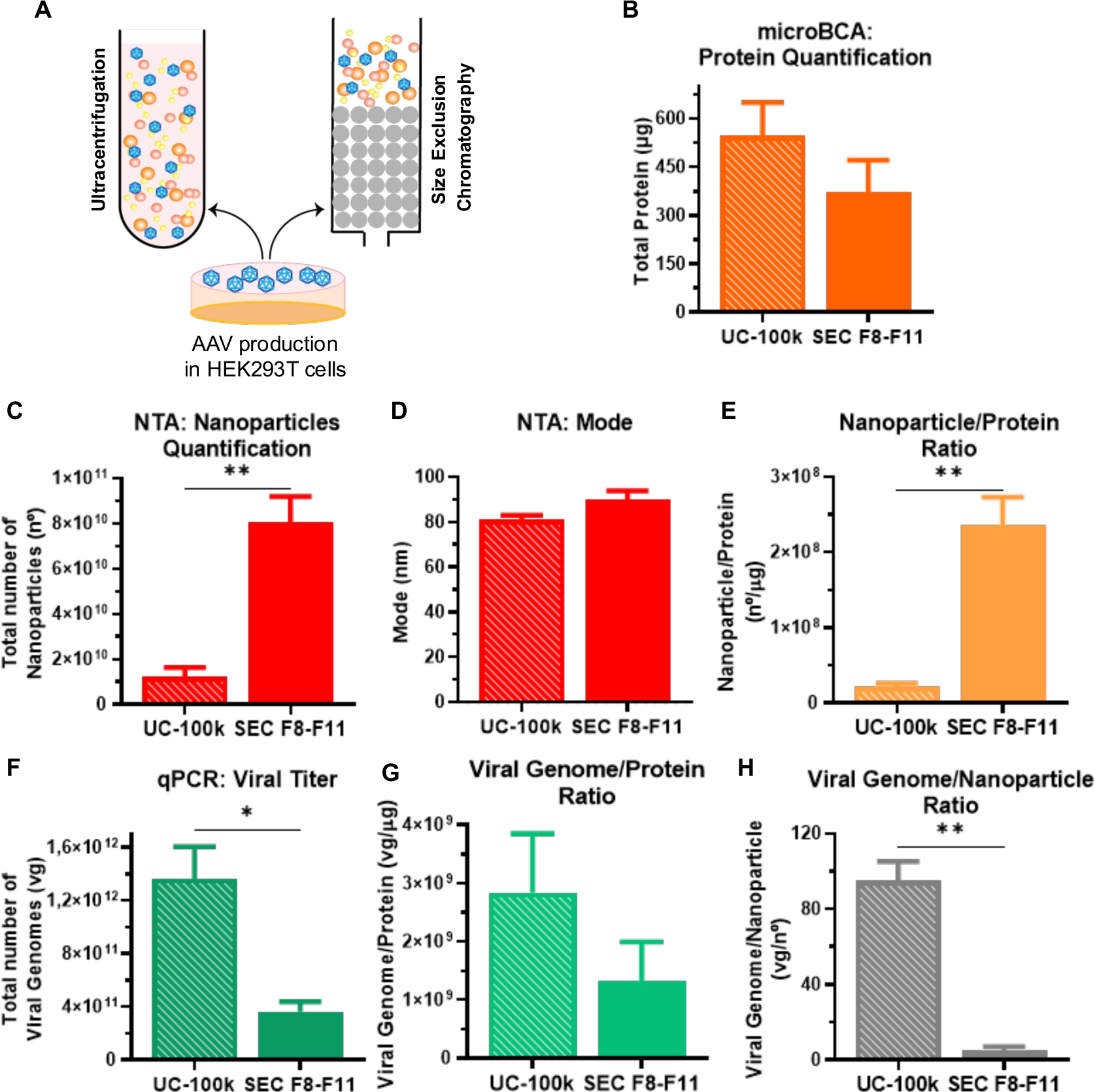
AAVs isolated by SEC contain more nanoparticles and less solo AAVs than EV-AAVs pelleted by UC-100k

No significant differences were observed in the total protein amount, nor in the size of the EV-AAVs isolated by the two methods (Figure 4.B and 4.D, respectively). Nevertheless, the yield of nanoparticles was drastically higher (6.6-fold) by SEC (Figure 4.C), suggesting a probable loss of EVs during UC. Nanoparticles per protein ratio was higher by SEC (Figure 4.E).

On the other hand, the viral titer was 3.8 times higher in EV-AAVs isolated by UC, which may indicate that UC simultaneously isolate AAVs, EVs and EV-AAVs. As result, the heterogeneous mixture of EV-AAVs isolated by UC present a higher viral genome to nanoparticle ratio (SEC: 5.023 ± 1.879; UC-100k: 94.77 ± 10.38) when compared to EV-AAVs isolated by SEC (Figure 4.G).

To compare the *in vitro* transduction of EV-AAVs isolated by UC and SEC, N2a cells were incubated with 10 000 viral genomes/cell. At 48 h post-infection, the percentage of cells expressing GFP was determined by FACS (Figure 5 and Supplementary Figure 5). The highest percentage of GFP-positive cells was detected in cells treated with EV-AAVs isolated by SEC (8.9 ± 0.75) (Figure 5.B-D, Supplementary Figure 5). Interestingly, EV-AAVs isolated by SEC or UC showed higher N2a cells transduction than solo AAVs (Figure 5.B-D).

**Figure 5-.**
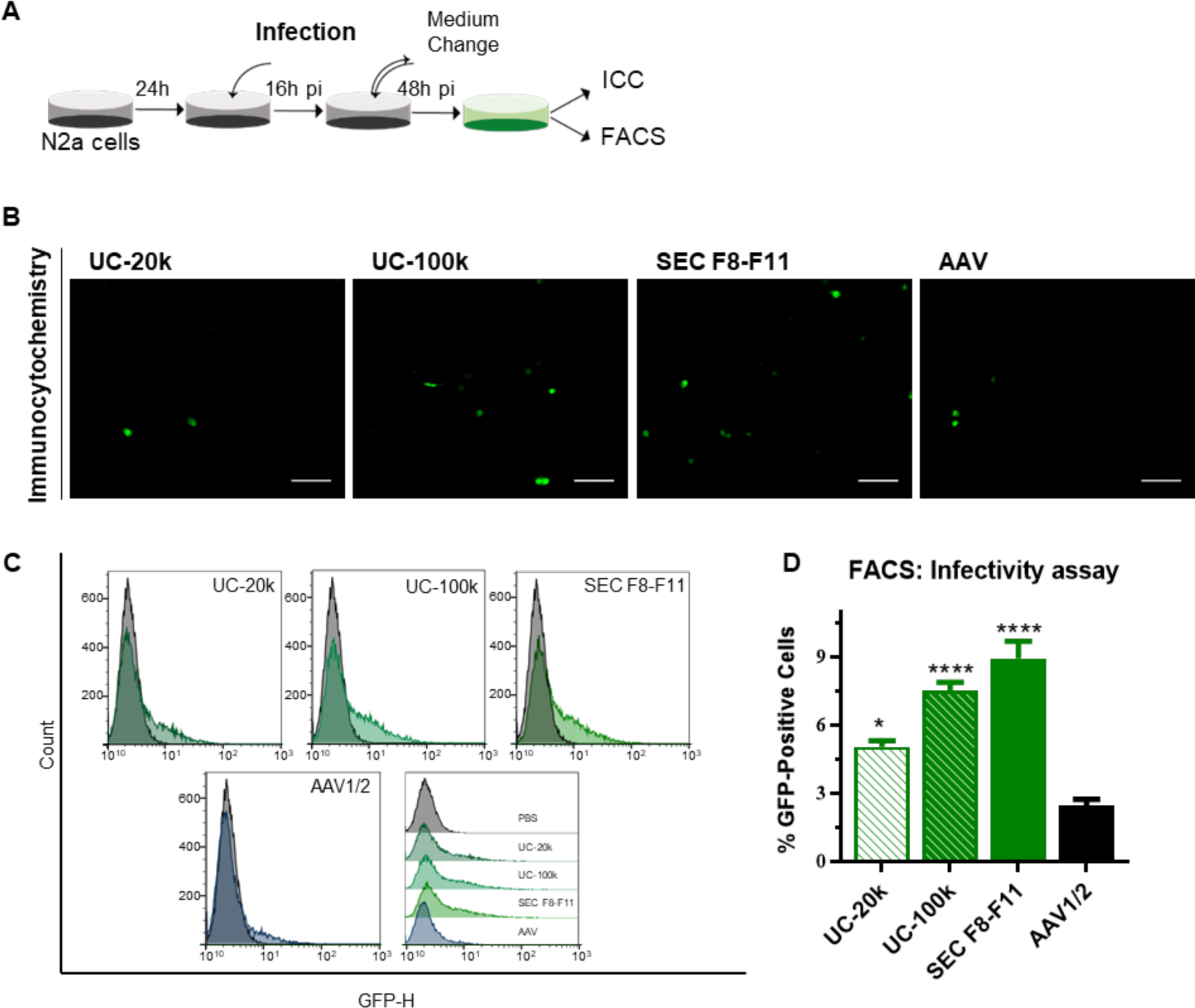
EV-AAVs mediate higher transduction efficiency than solo AAVs, particularly when purified by SEC

Since EV-AAVs isolated by SEC contained higher yields of nanoparticles, with less co-isolated contaminant proteins and solo AAVs, we proceeded to *in vivo* experiments with EV-AAVs isolated by SEC (F8-F11).

### 2.3. EV-AAVs transduce C57BL/6 mice brains upon intravenous injection

Given the *in vitro* efficacy of EV-AAVs purified by SEC, we next evaluated the possibility of using EV-AAVs as a non-invasive delivery system for the treatment of CNS diseases. For this purpose, WT C57BL/6 neonatal mice (P1) were intravenously injected in the facial vein with 1×10^10^ vg of EV-AAVs expressing GFP (Figure 6.A). Mice were sacrificed at P50 or P100 and immunohistochemistry against GFP was performed in sagittal brain sections to evaluate neuronal transduction.

**Figure 6 -.**
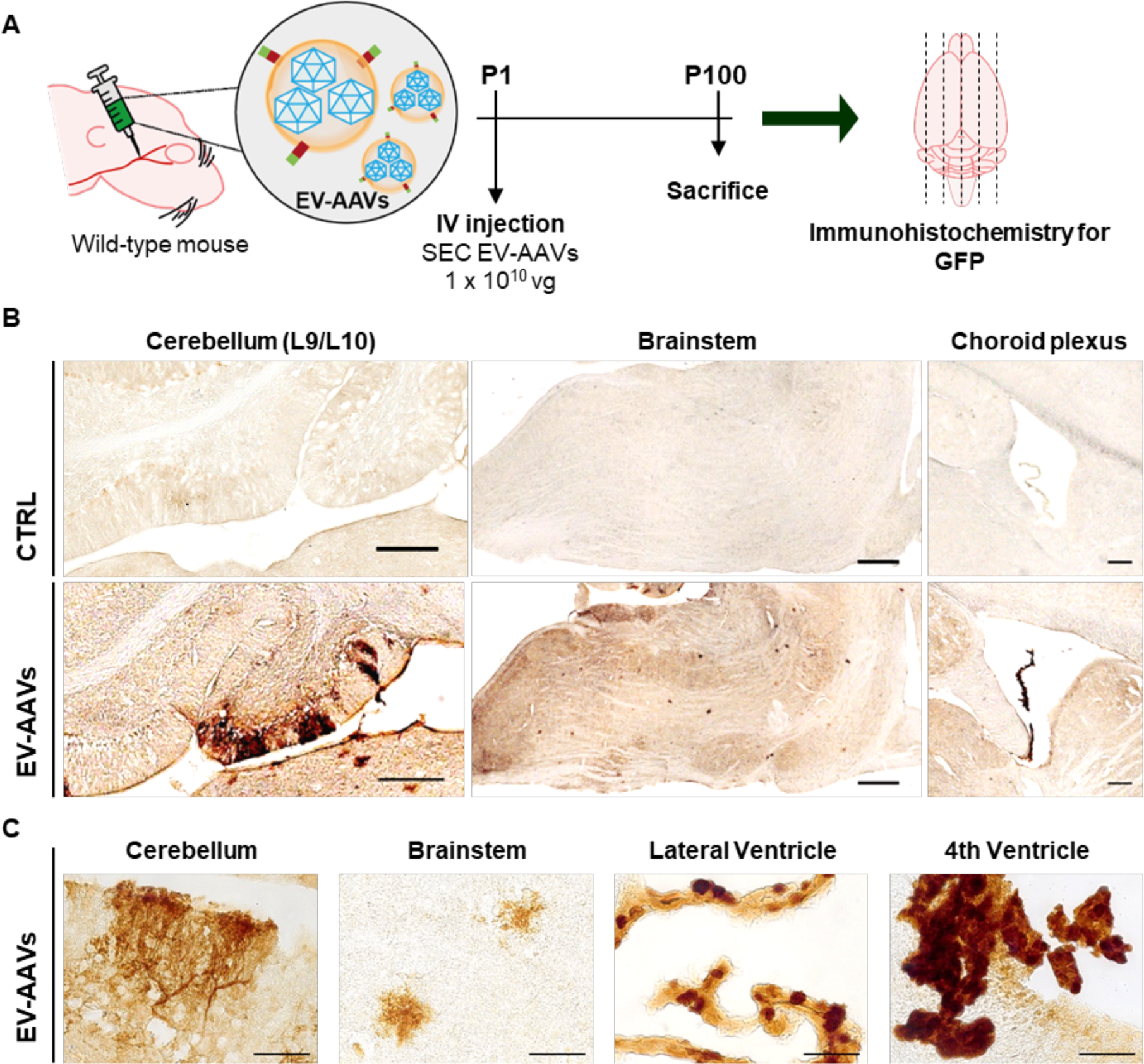
Intravenously injected SEC EV-AAVs are able to cross the BBB and transduce wild-type mouse brains

GFP expression was mostly observed in the cerebellum (particularly in Purkinje cells at lobule 10), brainstem and ependymal cells of the choroid plexus (lateral and fourth ventricles) (Figure 6.B-C). GFP expression was similar between animals sacrificed at P50 or P100. These results show that biologically formed EV-AAVs targeting the brain are able to cross the BBB and transduce the CNS.

## 3. Discussion

Adeno-associated viral (AAV) vectors are one of the preferential vehicles for gene therapy targeting the central nervous system (CNS)^3^. AAVs can infect both dividing and non-dividing cells, including neurons. Additionally, they exhibit no cytotoxicity and, in comparation with other viral vectors, have mild immunogenicity, allowing high efficiency and sustained expression of transgenes^29–31^. However, the use of AAVs has some limitations, such as the small transgene size capacity, the presence of biological barriers that limit gene delivery, such as the BBB, or the induction of an immune response due to capsid similarities with wild-type AAVs. To overcome these challenges, AAV capsids and genomes have been extensively engineered^10^.

In 2012, it was demonstrated that, during a normal production of AAVs in HEK293T cells, AAVs are also secreted in association with EVs^13^. EVs have been largely studied due to their unique characteristics and possible application, for example, as gene delivery vectors^32^. EVs are naturally formed vesicles, with a lipid membrane identical to the cell membrane, involved in intercellular communication. These small vesicles can travel through circulation, cross biological barriers, such as BBB and deliver their cargo to deep tissues^33–35^. Similar to EVs, EV-AAVs are also able to cross the BBB upon systemic injection and transduce neuronal cells, with the same tropism as standard AAVs, but with higher efficiency at lower vector doses^13, 18^. Moreover, it was observed that EV-AAVs transduce cells more efficiently than solo AAVs both *in vitro* and *in vivo*^14, 15, 16–19, 28, 36–38, 39^. Despite the relevance of this discovery, to date, only AAV serotypes able to cross the BBB (e.g. AAV9 and AAV8) have been used to target the CNS *in vivo*, which does not eliminate the possible contribution of solo AAVs to these observations^14, 18, 20^. Moreover, in these studies the purification method used for the isolation of EV-AAVs was based on UC^13, 18^. Although UC has been the most used protocol for the isolation of EV and EV-AAVs, solo AAVs can be co-isolated with EV-AAVs upon UC isolation^14^. Moreover, UC is limited by the lack of reproducibility, purity, and scalability which hampers its translation into the clinical setting. In this context, there was a need to optimize a reproducible and scalable protocol for the isolation of pure and biologically formed EV-AAVs, translatable for future clinical applications.

Previously, our group optimized a protocol for the isolation of EVs from human plasma by SEC, a method that renders highly pure EVs, without compromising their integrity and functionality^21, 22^. Here, we adapted this protocol to isolate EVs from conditioned media of AAV-producing HEK293T cells. To understand in which SEC F(s) EV-AAVs were isolated, we characterized the first 30 fractions of 500 uL eluted from the column. A large amount of proteins was obtained between F12 and F26, which we considered to be “free protein”^40^. Pure EVs were present mostly between F8 and F11, as shown by the presence of flotillin 1 (EVs marker) and absence of calnexin (cellular marker). Markers of AAV capsids (VP1-3) were also detected in these fractions. Since the pore size of the chosen SEC column is 70 nm, solo AAVs (∼ 25 nm) are not expected to elute in the same fractions of EVs. This indicates that AAVs might be eluted in association with EVs. Indeed, the fractions containing EV-AAVs (F7 to F11) showed to be more biologically active than those containing solo AAVs (i.e. F12, F13), as shown by the higher expression of GFP in F7 to F11 upon infection. Our results suggest that it is possible to isolate potent and naturally formed EV-AAVs, with less contaminant proteins and solo AAVs, by SEC.

Following the optimization of the SEC protocol, we next compared EV-AAVs isolated by UC-100k and SEC. Both methods isolated EVs with a similar size, however SEC allowed a yield 6.6-fold higher compared to UC, suggesting that SEC is more efficient in the recovery of EVs from the media. On the other hand, viral titer was 3.8-fold higher on EV-AAVs isolated by UC, which might be explained due to higher co-isolation of solo AAVs with EV-AAVs by UC. In fact, the centrifugation speed used is close to the sedimentation coefficient of AAVs^41^.

Based on that, the differences observed in the viral genome/nanoparticle ratio in SEC and UC may be attributed to not only a higher number of EVs without AAVs isolated by SEC, but also to the co-isolation of solo AAVs with EVs during UC. Immunoaffinity-based methodologies can be further used in conjunction with SEC to specifically isolate an EV-AAV subpopulation, further increasing the homogeneity of the sample^42^. Additionally, EV-AAVs isolated by UC were 10.88-fold less pure than those isolated by SEC, according to the Nanoparticles/Protein ratio previously described by Webber and Clayton^43^.

In any case, EV-AAVs isolated by UC and SEC were more potent at transduction than solo AAVs *in vitro*, possibly due to the higher neuronal targeting capacity of EV-AAVs, as compared to solo AAVs, due to the presence of RVg peptide on their surface^14, 44^.

Having demonstrated the efficiency and advantages of isolating EV-AAVs by SEC *in vitro*, we aimed at evaluating the possibility of using EV-AAVs as a non-invasive delivery system targeting the CNS. To determine whether EV-AAVs could cross the BBB without the contribution of AAV serotype, we chose mosaic AAV1/2 vectors for the generation of EV-AAVs. Mosaic AAV vectors combine the properties of both parental AAVs, including neurotropic features; however, they are still unable to efficiently cross the BBB on their own^25^. To direct EV-AAVs to the brain, the RVg peptide was engineered on the surface of EVs, as previously demonstrated by György and colleagues^14^. RVg peptides bind to acetylcholine receptors and selectively target neuronal cells and brain endothelial cells, enabling EVs to cross the BBB^44, 45, 47^.

Following an intravenous injection of SEC EV-AAVs in neonatal mice, we showed that EV-AAV1/2 coated with RVg can cross the BBB and transduce mouse brains. To our knowledge, this is the first time that EV-AAVs comprising AAV serotypes without capacity for crossing the BBB were used. This proves not only that EV-AAVs are naturally formed, but also that they are capable to cross barriers, such as the BBB and the blood-cerebrospinal fluid barrier (BCSFB), without AAVs serotype contribution.

Interestingly, the high rate of transduction of ependymal cells makes EV-AAVs strong candidates to gene delivery through the cerebrospinal fluid (CSF), allowing a long-term supply of transgenic product^46^. On the other hand, the capacity to target the cerebellum in mice highlights the potential of EV-AAVs as a new gene delivery platform for diseases such as spinocerebellar ataxias that mainly affect the cerebellum^48^.

Finally, it is important to note that in the present work, we used 10 to 38-fold fewer viral genomes for IV injection (1×10^10^ vg/animal) compared to what has been used with solo AAV9 (1×10^11^ - 3.8×10^11^ vg/animal)^3, 49–52^. Based on this, we can speculate that using a higher amount of EV-AAVs, but still smaller than solo AAV9, it would be possible to achieve a global CNS transduction.

## 4. Limitations

In both studies investigating the transduction of EV-AAVs in N2a cells, it was not possible to determine the specific impact of cDNA inside vesicles, but not packaged within the AAV capsid. To assess the individual contribution of the cDNA to transduction, it would be necessary to repeat the EV production protocol without using the “AAV rep and cap genes” plasmid. Earlier work has suggested that transgene DNA outside of the capsid is unlikely to play a role in transduction^13^. Furthermore, plasmid DNA loaded into EVs is not efficient at transgene expression^53^. However, it is evident that the presence of AAVs associated with EVs is the primary factor affecting transduction efficiency, as demonstrated by the observed brain transduction in rodents 100 days after the injection of EV-AAVs.

## 5. Conclusions

In conclusion, SEC allows the recovery of pure and biologically formed EV-AAVs that can be scaled up for future clinical applications. Given their outstanding features, not only at evading the immune system and achieving higher cell transduction but also at crossing biological barriers, such as the BBB, EV-AAVs may be a promising approach for gene therapy of CNS disorders.

## 6. Materials and Methods

### *In vitro* experiments

All work involving cells and viral vectors was performed in dedicated biosafety cabinets and incubators separated from those used for maintaining cell lines. The equipment and reagents in contact with cells in culture and viruses were sterilized and materials virally contaminated were properly rinsed or disposed, according to good laboratory practices.

#### 6.1. Cell lines

Human Embryonic Kidney cells 293T (HEK293T) and Mouse Neural Crest-derived cell line (Neuro 2A cells) were obtained from the American Type Culture Collection cell biology bank (CRL-11268 and CCL-131, respectively). Both cell lines were cultured in Dulbecco’s Modified Eagle’s Medium (DMEM) supplemented with 10 % Fetal Bovine Serum (FBS) and 1 % Penicillin-Streptomycin (P/S). Cells were maintained in 5 % CO_2_ at 37°C, unless mentioned otherwise. Sterile Phosphate Buffered Saline (PBS), pH = 7.4, and 0.05 % trypsin were used for cell passage.

#### 6.2. Production of AAVs and EV-AAVs

Twenty four hours before transfection, 1.05 x 10^7^ HEK293T cells were plated into 15-cm dishes and cultured in DMEM supplemented with 10 % FBS, 1 % P/S and 1M hydroxyethyl-piperazine ethane sulfonic acid (HEPES) buffer. AAVs were produced according to the standard transfection method^26^. Briefly, four plasmids were used: i) a plasmid encoding the enhanced green fluorescent reporter gene (GFP) under the chicken beta-actin (CBA) promoter placed between the ITRs (pITR) sequences, ii) two plasmids containing the wild-type (WT) AAV genome *rep* and *cap* ORFs (AAV1 and AAV2) and iii) a plasmid encoding the adenovirus proteins (E1A, E1B, E4 and E2A) along with the adenovirus’ RNAs necessary for helper functions (pHelper). A fifth plasmid encoding for the Rabies virus glycoprotein peptide-platelet-derived growth factor receptor (RVg-PDGFR) transmembrane domain was also transfected^14^.For an efficient transfection, Polyethylenimine (PEI) linear MW 40000 (Polysciences, Inc) was used as the transfection reagent.

Medium was replaced by fresh DMEM 2 % EV-free FBS, 1 % P/S and 1 % HEPES 16h post-transfection to reduce bovine EVs.

Conditioned medium was collected at 48 h and 72 h post-transfection and centrifuged for 10 min at 500 g followed by another centrifugation at 1 000 g for 15 min to remove cell debris. Medium was stored at −80°C. Cells were harvested for AAV purification.

#### 6.3. Isolation of EV and EV-AAVs by Differential Centrifugation

Differential centrifugation was performed using a 70 Ti rotor in an Optima XE-100 ultracentrifuge (Beckman Coulter). Conditioned medium was centrifuged for 60 min at 20 000 g and supernatant was centrifuged once again at 100 000 g for 90 min. Resulting pellets were resuspended in PBS.

#### 6.4. Isolation of EV-AAVs by Size Exclusion Chromatography

Conditioned medium was concentrated using centrifugal concentrators (Vivaspin 20/Vivacell 70, Sartorius) or pressure concentrators (Vivacell 250, Sartorius) with a MWCO of 10 000 kDa. For EV-AAV purification, commercial agarose-based size exclusion columns (qEVoriginal, Izon Science) were used. For each assay, 1 mL of concentrated medium was loaded on qEVoriginal columns followed by PBS. Fractions of 0.5 mL were collected and immediately used or stored at −80°C before further analysis.

For *in vivo* experiments, after SEC, an additional concentration step was performed using centrifugal concentrators (Vivacell20, Sartorius).

#### 6.5. Isolation of AAVs

AAVs were purified based on a Fast Protein Liquid Chromatography (FPLC) in our core facility ViralVector (CNC), using the AKTA pure 25 system (GE Healthcare, Life sciences). Purified rAAVs were subsequently concentrated using centrifugal concentrators (Amicon Ultra-15 and Ultra-0.5, Merck) with a molecular weight cut-off (MWCO) of 100 000 kDa and supplemented with 0.001 % Pluronic F-68 100X (Life Technologies) to avoid aggregation.

#### 6.6. Protein quantification

Total protein was quantified in samples isolated by UC and SEC through bicinchoninic acid (BCA) (Thermo Scientific), using lysed (described below), and MicroBCA (Thermo Scientific) assays, using non-lysed samples.

#### 6.7. Western Blot

All samples used for western blot analysis, were lysed with RIPA buffer [150 mM NaCl, 50 mM Tris-Cl, 5 mM ethylene glycol tetraacetic acid (EGTA), 1 % Triton X-100, 0.5 % deoxycholic acid (DOC), 0.1 % sodium doceylsulphate (SDS) pH = 7.5) supplemented with a protease inhibitor cocktail (Roche) and 200 mM phenylmethylsulphonyl fluoride (PMSF), 1 mM dithiothreitol (DTT), 1 mM activated sodium orthovanadate and 10 mM sodium fluoride (NaF)], with the exception of solo AAVs purified by FPLC. Protein extracts from HEK293T cells were also used as reference. Protein extracts were resolved in SDS-polyacrylamide gels (4 % stacking and 10 % running) and transferred onto polyvinylidene difluoride (PVDF) membranes (Immobilon-P, EMD Millipore), according to standard protocols. After protein eletrotransfer, membranes were stained with Ponceau S to visualize total protein and subsequently washed with 0.1 M NaOH. Then, membranes were blocked by incubation in 5 % non-fat milk powder in 0.1 % Tween 20 in Tris buffered saline (TBS-T), for 1 h at room temperature (RT). Immunoblotting was performed overnight at 4 °C with the following primary antibodies: anti-calnexin H70 (rabbit polyclonal, Santa Cruz Biotechnology, 1:1000), anti-flotilin-1 clone 18 (mouse monoclonal, BD transduction laboratories, 1:400), anti-AAV (VP1/VP2/VP3) clone B1 (mouse monoclonal, American Research Products, Inc, 1:1000), followed by incubation with corresponding alkaline phosphatase-coupled secondary antibodies (goat anti-rabbit polyclonal antibody, Thermo Scientific, 1:10000; or goat anti-mouse polyclonal antibody, Invitrogen, 1:10000). The presence of the antigens of interest was observed with Enhanced Chemifluorescence substrate (ECF, Ge Healthcare) and chemifluorescence imaging (ChemiDoc^TM^ Imaging System, BioRad).

#### 6.8. Quantification of AAVs by Real-Time PCR

Absolute quantification of rAAV titer (vg/µL) was performed by quantitative real-time polymerase chain reaction (PCR) using the AAVpro™ Titration Kit (for Real Time PCR) Ver.2 (Takara Bio Inc), according to the manufacturer’s instructions. Absolute quantification was based on the amplification of the inverted terminal repeats (ITRs) of AAVs using the standard curve method.

#### 6.9. Nanoparticle Tracking Analysis (NTA)

Measurements of nanoparticle size and concentration in EV-AAVs preparations were performed by Nanoparticle Tracking Analysis (NTA). A NanoSight NS300 instrument (Marvell Panalytical, Malvern) with a 488 nm laser and sCMOS camera module (Malvern Panalytical, Malvern, United Kingdom) was used following the manufacturer’s instructions. Five videos of 30 s were recorded for each sample using a syringe pump speed of 40. Measurements were performed using NTA 3.2 software (Malvern).

#### 6.10. Transmission Electron Microscopy (TEM)

Transmission electron microscopy was performed according to Thierry *et al.*, 2006 ^54^. Briefly, EV-AAVs were fixed with 2% paraformaldehyde (PFA) and deposited on Formvar-carbon coated grids (TAAB Laboratories Equipment) for 20 min. Grids were washed with PBS and fixed for 5min with 1% glutaraldehyde. Following a cycle of washes using distilled water, grids were contrasted with a uranyl-oxalate solution (pH = 7) for 5 min and transferred to methyl-cellulose-uranyl acetate for 10 min on ice. Images were obtained using a Tecnai G2 Spirit BioTWIN electron microscope (FEI) at 80 kV.

#### 6.11. *In vitro* transduction Assay

To evaluate the transduction efficiency of EV-AAVs and EV + AAV, Neuro 2A (N2a) cells were seeded in either 24-well plates (5×10^4^ cells/well) or 48-well plates (2×10^4^ cells/well), for fluorescence-activated cell sorting (FACS) and immunocytochemistry (ICC) analysis, respectively. On the following day, half of the medium was collected and cells were infected at 10 000 vg/cell, with different samples equally diluted in PBS. Sixteen-hours post-vector addition, the medium was replaced. Transgene expression was evaluated 48 hours post-infection.

#### 6.12. Fluorescence-Activated Cell Sorting (FACS)

Two days post-infection, infected N2a cells were processed for flow cytometry analysis. Briefly, cells were harvested, washed with cold PBS, collected by centrifugation and resuspended in PBS in conic tubes (BD, Biosciences). Samples were kept on ice before being analyzed for GFP expression in a FACS Calibur flow cytometer (BD, Biosciences). A total of 2×10^4^ cells were scanned during each acquisition and GFP fluorescence was evaluated in the FL-1 channel. Data analysis was performed using FlowJo software (Tree Star Inc., Ashland, USA).

#### 6.13. Immunocytochemistry

Two days post-infection, infected N2a cells were washed with PBS and fixed in 4 % paraformaldehyde in PBS, for 15 min at RT. After fixation, cells were washed and permeabilized with 0.1 % Triton X-100/PBS, for 5 min at RT. Blockage of non-specific staining was performed with 10 % bovine serum albumin (BSA) in PBS, for 30 min at 37 °C, and cells were then incubated overnight at 4 °C with the rabbit anti-GFP primary antibody (1:1000, Invitrogen) diluted in 3 % BSA in PBS. Subsequently, cells were washed and incubated for 45 min at 37 °C with the goat anti-rabbit secondary antibody (1:250, Thermo Scientific), also diluted in 3% BSA in PBS.

Images were obtained using CELENA^®^ S digital cell imaging system (Logos Biosystems) with a 10X objective.

### *In vivo* Experiments

#### 6.14. Animals

Neonatal (P1) WT (N = 7) pups (C57BL/6 background) were used to evaluate the transduction efficiency of the EV-AAV purified by SEC, upon a minimally invasive intravenous injection.

Animals were housed in a temperature-controlled room maintained on a 12-hour light / 12-hour dark cycle. Food and water were provided *ad libitum*. Experiments involving mice were previously approved by the Responsible Organization for the Animals Welfare (ORBEA) of the Faculty of Medicine, Directorate-general for Food and Veterinary Medicine (DGAV) and the Center for Neuroscience and Cell Biology of the University of Coimbra (FMUC/CNC, Coimbra, Portugal). All experiments were carried out in accordance with the European Union Community directive (2010/63/EU) for the care and use of laboratory animals, transposed into the Portuguese law in 2013 (Decree Law 113/2013). The involved researchers received adequate training (Felasa-certified course), and certification to perform animal experiments from the Portuguese authorities (Direcção Geral de Alimentação e Veterinária, Lisbon, Portugal). All efforts were made to minimize animal suffering.

#### 6.15. Neonatal Intravenous Injections

C57BL/6 pregnant mice were housed and monitored daily from embryonic day 17 to 21, with the least possible disturbance, to ensure that newborn pups could be dosed with vectors on P1.

Newborn mice were initially rested on a bed of ice for approximately 1 min for anesthetization. EV-AAV samples, isolated by SEC, containing a total of 1×10^10^ vg, diluted in 50 μL of PBS, were manually injected into the facial vein of seven newborn mice using a 100 µL Hamilton syringe connected to a 30-gauge beveled tip needle (Hamilton). A correct injection was verified by noting blanching of the vein. After injection, pups were identified by toe tattooing, carefully cleaned, rubbed with their original bedding to prevent rejection by the mother, and then returned to their original cage. Animals were sacrificed 50 (N = 3) or 100 (N = 4) days post-injection, for histological analysis of the brain.

### Histological Processing

#### 6.16. Tissue Collection and Preparation

Mice were terminally anesthetized through the intraperitoneal route and transcardially perfused with cold PBS (pH = 7.4). Brains were excised and post-fixed with 4 % PFA for 48 h at 4 °C, and then transferred to 20 % sucrose in PBS for cryoprotection. Brains were frozen and stored at −80 °C upon sinking (approximately 48h later).

Sagittal sections were subsequently cut at 30 μm thickness, using a cryostat (CRYOSTAR NX50, Thermo Scientific) at −20 °C. Sections were stored in 48-well trays as free-floating sections in PBS supplemented with 5 % sodium azide. Trays were stored at 4 °C until further processing.

#### 6.17. Bright-field immunohistochemistry

After being rinsed in PBS, free-floating sections were incubated in 0.1 % phenylhydrazine in PBS (Merck) for 30 min at 37 °C, to block endogenous peroxidases. Tissue blocking and permeabilization was subsequently performed in blocking solution (0.1 % Triton X-100 containing 10 % Normal Goat Serum (NGS, Gibco) in PBS), for 1 h at RT. Brain slices were then incubated overnight at 4 °C with the rabbit anti-GFP primary antibody (1:1000, Invitrogen) diluted in blocking solution. Following three washing steps in PBS, free-floating sections were incubated for 2h at RT with the anti-rabbit biotinylated secondary antibody (Vector Laboratories) diluted in blocking solution (1:250). Subsequently, free-floating sections were rinsed and bound antibodies were visualized by the Avidin-Biotin complex (ABC) amplification system (Vectastain ABC kit, Vector Laboratories) using 3,3′-diaminobenzidine tetrahydrochloride (DAB Substrate Kit, Vector Laboratories) as substrate. The reaction was stopped by washing sections in PBS, after achieving optimal staining.

Sections were mounted on gelatin-coated slides, hydrated with ultrapure water, and dehydrated in an ascending ethanol sequence (70 %, 95 % and 100 %). Slides were then cleared with xylene solution and finally coverslipped with Eukit (O. Kindler GmbH & CO). Images were acquired using a Zeiss Axio Imager Z2 microscope (Carl Zeiss Microscopy GmbH), equipped with a High Resolution Colour Camera. Images of cerebellum, brainstem and choroid plexus were obtained with a Plan-Apochromat 20X/0.8 M27 or 63x/1.4 objective.

#### 6.18. Statistical Analysis

All statistical analyses were performed using the GraphPad Prism software (version 9.0.0). Data are presented as mean ± standard error of mean (SEM). Unpaired student’s t-tests and One-way ANOVA tests were performed when applicable. Significance was determined according to the following criteria: * p < 0.05, ** p < 0.01, *** p < 0.001, and **** p < 0.0001.

## Acknowledgments

This work was funded by the European Regional Development Fund (ERDF), through the Centro 2020 Regional Operational Program; through the COMPETE 2020 – Operational Programme for Competitiveness and Internationalization, and Portuguese national funds via FCT under the projects: UIDB/04539/2020, UIDP/04539/2020, LA/P/0058/2020, SpreadSilencing (POCI-01-0145-FEDER-029716), ViraVector (CENTRO-01-0145-FEDER-022095), Fighting Sars-CoV-2 (CENTRO-01-01D2-FEDER-000002), BDforMJD (CENTRO-01-0145-FEDER-181240), ModelPolyQ2.0 (CENTRO-01-0145-FEDER-181258), and MJDEDIT (CENTRO-01-0145-FEDER-181266); ARDAT under the IMI2 JU grant agreement no. 945473 supported by the European Union’s H2020 programme and EFPIA; by the American Portuguese Biomedical Research Fund (APBRF) and the Richard Chin and Lily Lock Machado-Joseph Disease Research.

C.H. was supported by 2021.06939.BD; M.M.L. was supported by 2021.05776.BD; D.R.R. was supported by SFRH/BD/132618/2017; D.L. was supported by 2020.09668.BD; K.L. was supported by SFRH/BD/09513/2020; and A.C.S. was supported by 2020.07721.BD.

We thank M. Zuzarte (Faculty of Medicine, University of Coimbra) for electron microscopy imaging. We thank Luisa Cortes, Tatiana Catarino and Margarida Caldeira, the CNC MICC team for assistance with microscopy imaging. We thank all members of the L.P.d A. lab for all the support, discussion and comments.

## Author Contributions

C.H., M.M.L., M.S., L.P.d.A. and R.J.N. conceived and designed the experiments. C.A.M. shared knowledge and material. C.H., M.M.L., P.A., D.R.R., L.S.G., D.L., K.L., A.C.S., R.B., S.D. and M.S. performed the experiments. C.H., M.S., R.J.N. analyzed the data. C.H., M.S., R.J.N. wrote the first draft of the paper. All the authors reviewed and edited the paper.

---

**Supplementary Figure 1-.**
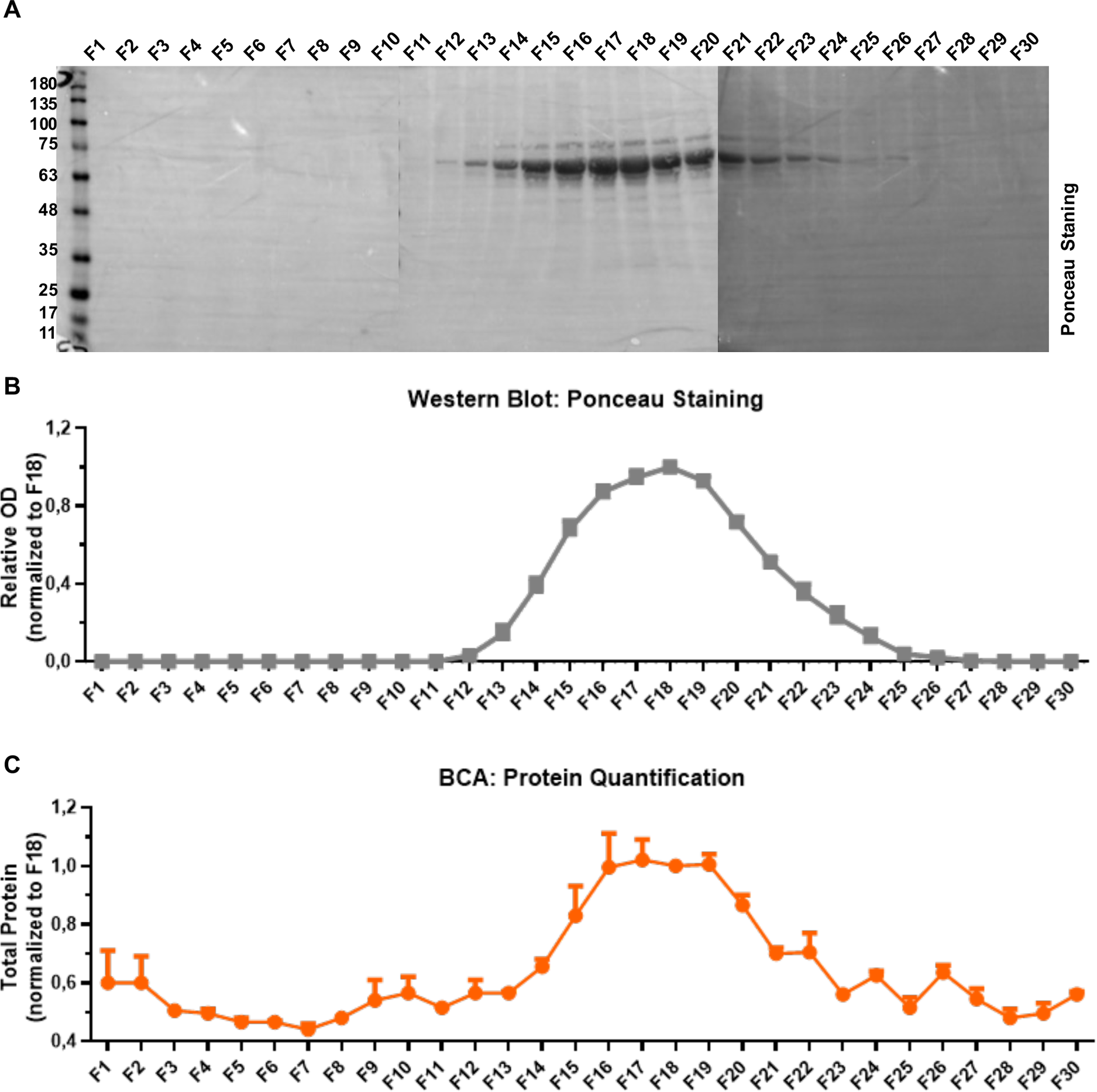
EV-enriched fractions present low protein contamination

**Supplementary Figure 2-.**
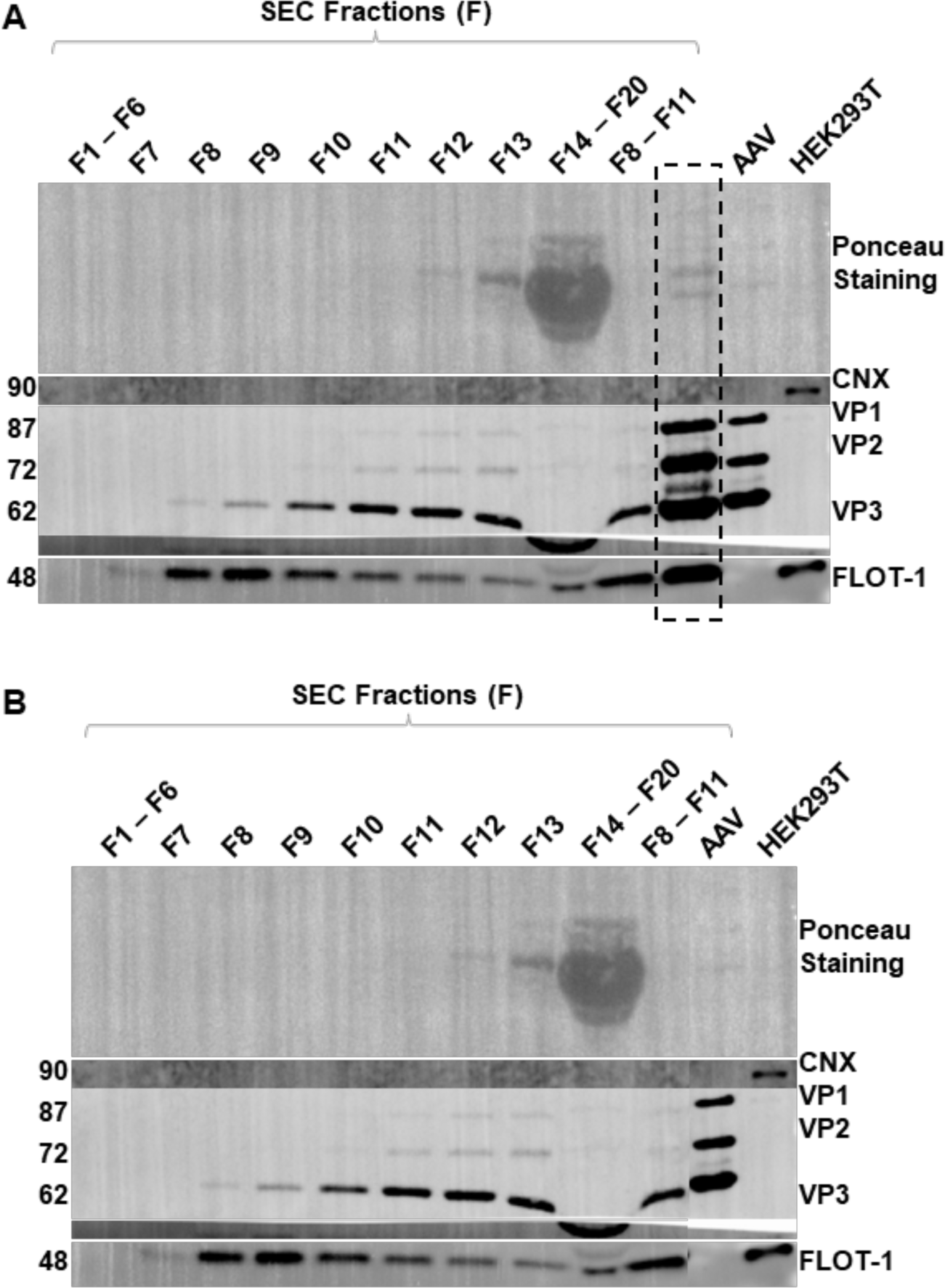
Uncropped western blot from main figure

**Supplementary Figure 3-.**
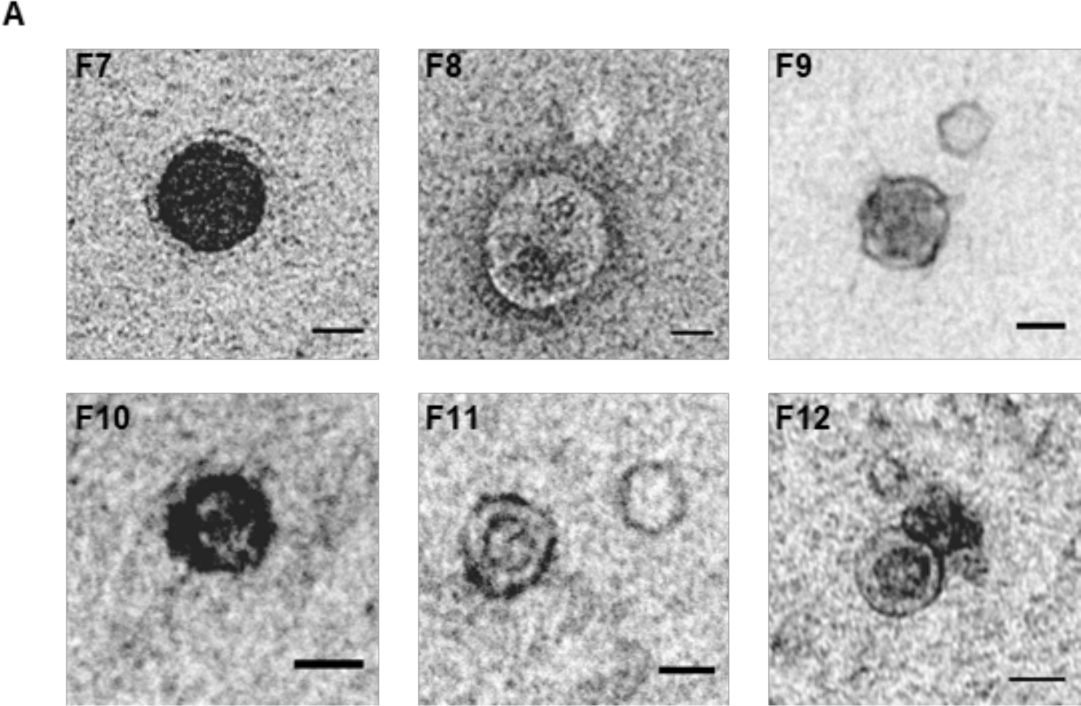
Characterization of SEC F7 to F12 by negative Transmission Electron Microscopy (TEM)

**Supplementary Figure 4-.**
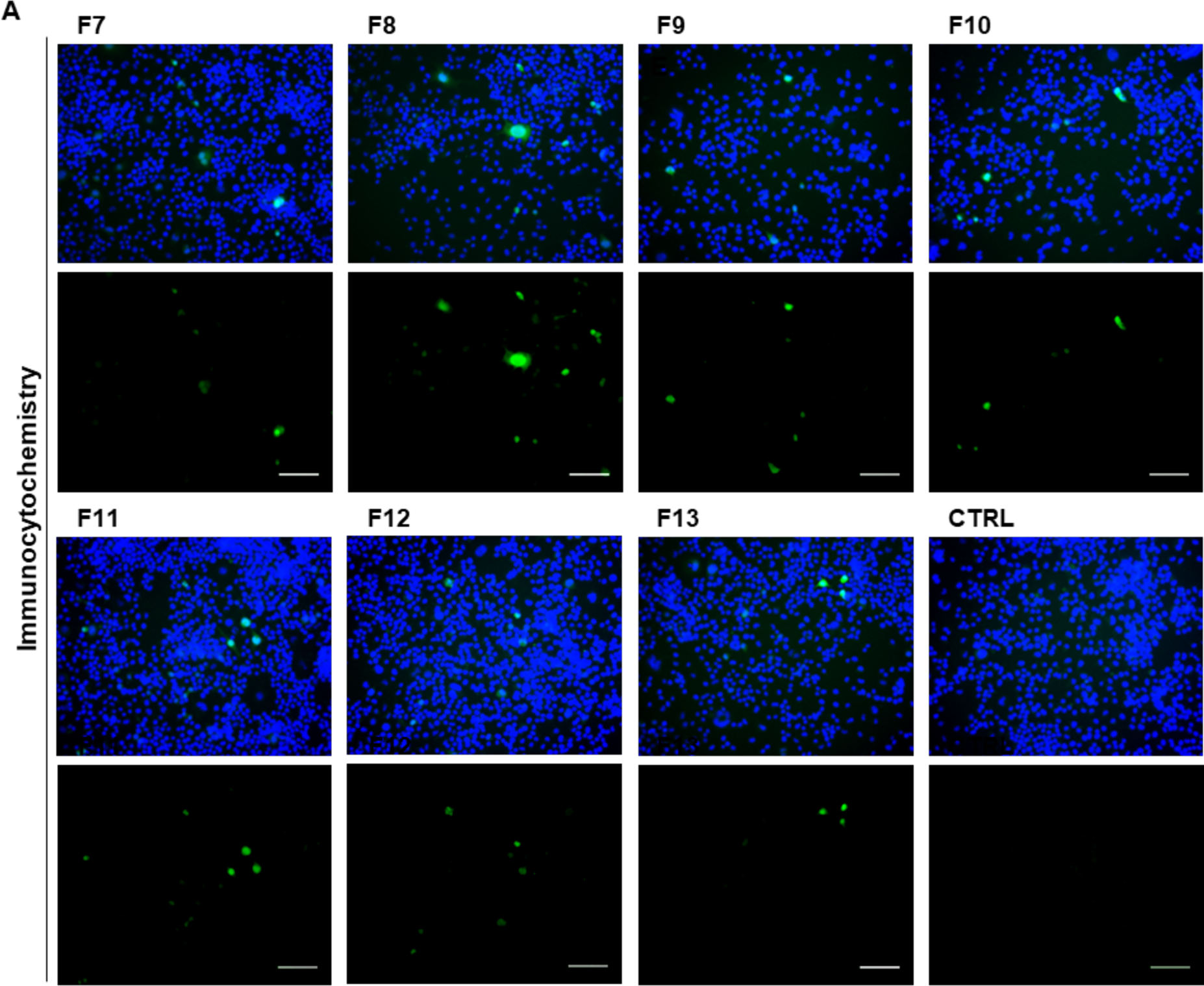
EV-AAVs (F7 to F11) were more efficient at transducing neuronal cells than solo AAVs (F13)

**Supplementary Figure 5-.**
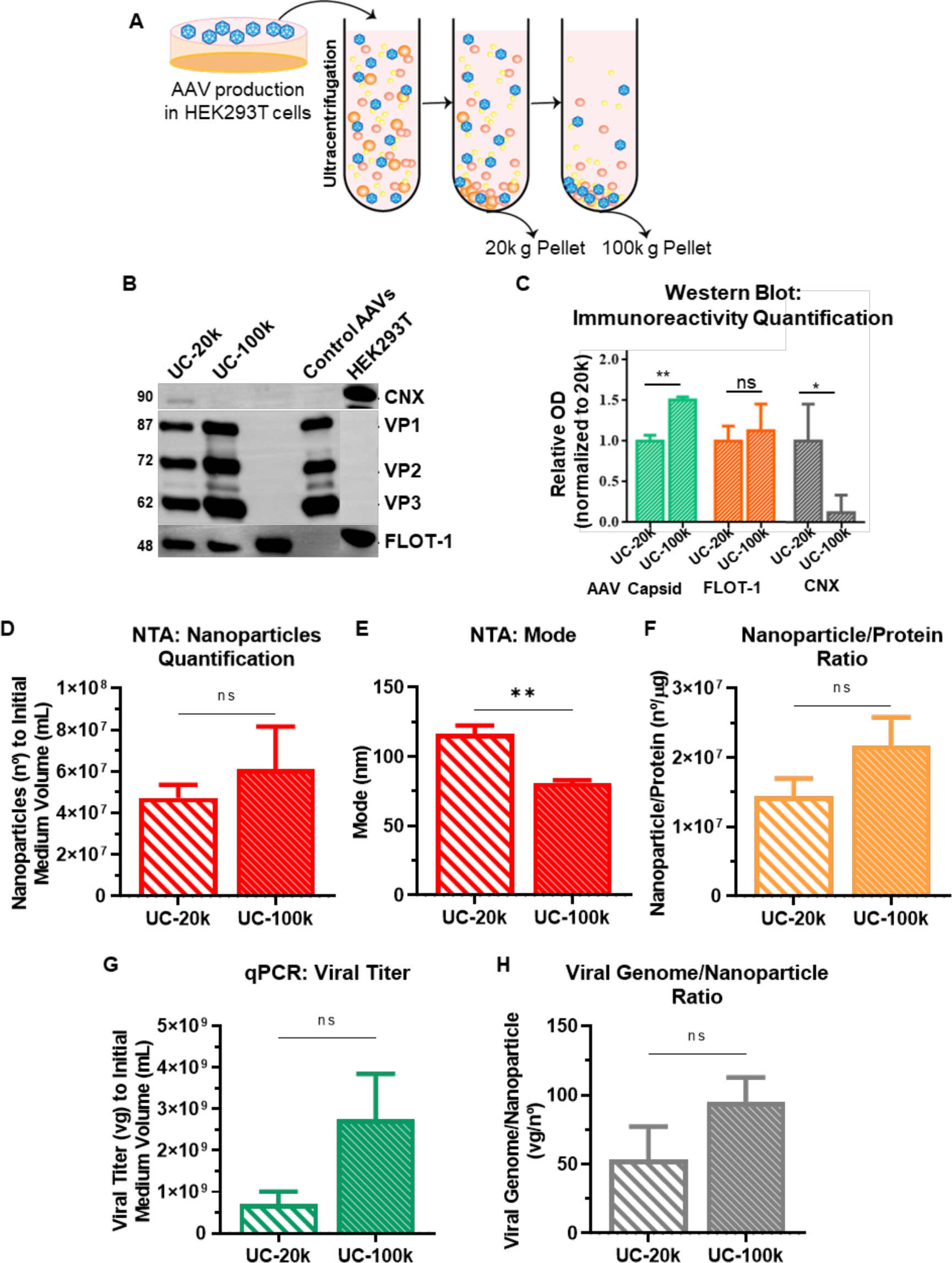
Two different populations of EVs are obtained during ultracentrifugation of AAV production medium at 20 000 g (UC-20k) and 100 000 g (UC-100k)

**Supplementary Figure 6-.**
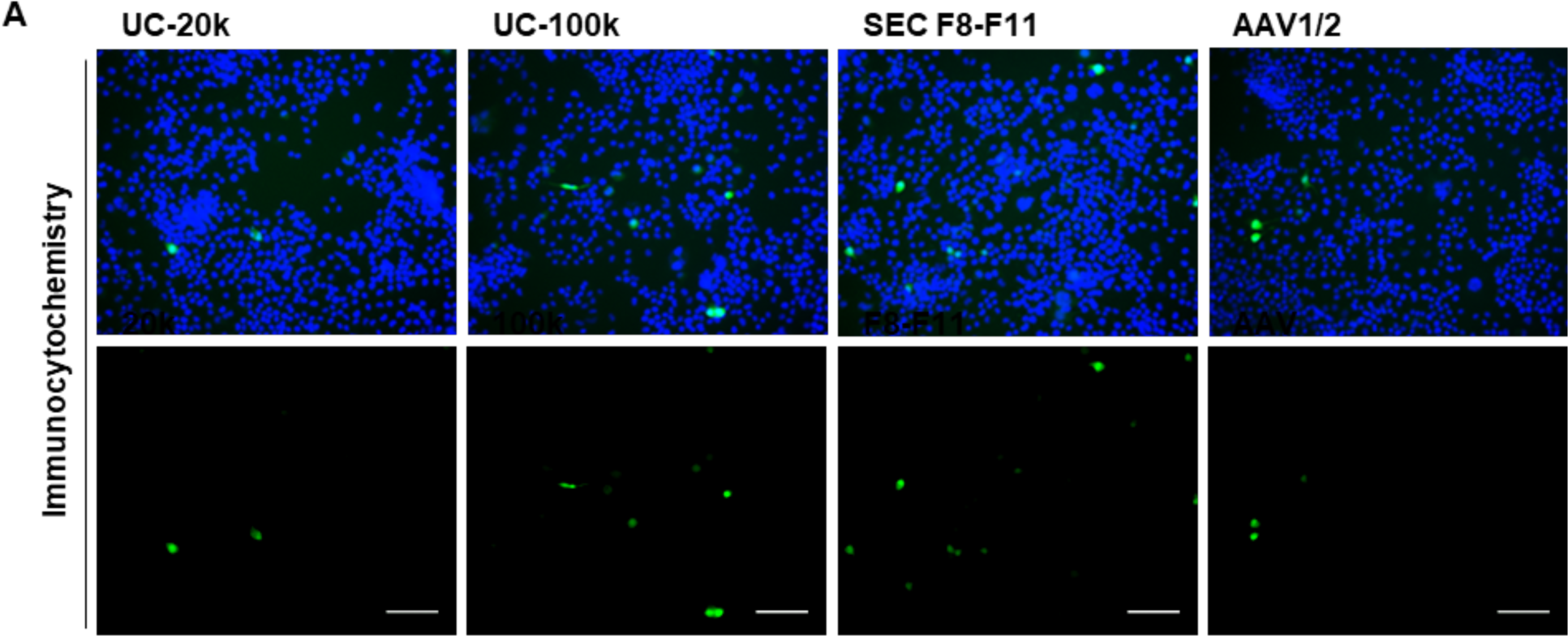
EV-AAVs are more efficient at transduction of cells than solo AAVs, particularly when purified by SEC

**FIGURE CAPTIONS:**

**Figure 7 - Workflow summarizing the protocol for the production and isolation of brain targeting EV-AAVs by SEC**

(A) EV-AAVs production using the standard protocol for AAV production. Briefly, HEK293T cells were transfected with four plasmids: one encoding the transgene of interest (GFP) between the ITRs (pITR), two plasmids containing the wild-type AAV 1 and 2 rep and cap ORFs (pAAVs) and a helper plasmid (pHelper). An additional plasmid encoding for the Rabies virus glycoprotein peptide (pBrain targeting peptide) was also transfected. In the following day, the culture medium was replaced with fresh medium supplemented with 2 % EV-free FBS. Conditioned medium was collected at 48h and 72h post-transfection. (B) The culture medium was then pre-cleared to remove cell debris by performing two low speed centrifugations or by using 0.22 µm filtration units. To concentrate the conditioned medium, 10 000 molecular weight cut-off ultrafiltration units were used. (C) One milliliter of concentrated medium was applied into the SEC column and 500 µL fractions were collected.

**Figure 8 – EV-AAVs can be separated by SEC**

(A) The presence of EV-marker FLOT-1, AAV capsid proteins (VP1, 2 and 3) and cell contamination contaminating protein (Calnexin, CNX) was analyzed by western blot. The same volume of fractions F1-F6, F7 to F13, F14-F20 and F8-F11 was loaded. Control AAVs purified by FPLC and HEK293T cell lysates were used as controls. (B) Relative expression of AAV capsid proteins (VP1, VP2, VP3) and FLOT-1, normalized to maximum OD, was quantified. AAV capsid proteins were detected in F8 and their presence increased in subsequent fractions. FLOT-1 was firstly detected in F7, with a higher expression between F8 and F11 and a subsequent decrease along following fractions. (C, D) Nanoparticle number and mode distribution determined by NTA. Nanoparticle concentration was higher in F8 and increased again from F10 to F13. (E) Nanoparticle size and the nanoparticle/protein ratio decreased along fractions. (F) Viral titer quantified by qPCR. Viral titer significantly increased along SEC fractions. (G) The number of viral genomes per protein increased between F8 and F10 and decreased from F11. (H) Number of viral genomes per nanoparticle augmented until F12. (B - H) Results are shown as mean ± SEM for F7-F13 (N = 3 per group, except in (B) – N = 6 per group, and (F) – N = 12 per group). See also Supplementary Fig. 1-3.

**Figure 9 - EV-AAVs (F7 to F11) are the most potent fractions in neuronal cells**

(A) Schematic representation of the transduction assay. Neuro 2A cells were infected individually at a MOI of 10 000 with SEC fractions F7 to F13. Non-transduced cells were used as control (CTRL). GFP expression was evaluated 48h post-infection by ICC (B) and measured FACS (C, D). F7 to F11 were the most potent fractions. (D) Statistical analysis relative to control was performed using Ordinary One-way ANOVA with Dunnett’s post-hoc test (* p < 0.05, ** p < 0.01, *** p < 0.001) and unpaired Student’s t-test was used to compare between fractions (# p < 0.05, ## p < 0.01). Results are shown as mean ± SEM for F7 to F13 (N = 4) and CTRL (N = 4). Scale bar = 200 µm. See also Supplementary Fig. 4.

**Figure 10 - AAVs isolated by SEC contain more nanoparticles and less solo AAVs than EV-AAVs pelleted by UC-100k**

(A) Schematic representation of the experimental design. EV-AAVs from AAV production medium were purified by UC and SEC in parallel. (B) Protein content measured by microBCA. (C, D, E) Nanoparticle number and mode distribution determined by NTA. The nanoparticle yield was significantly higher by SEC. The nanoparticle size mode was similar between UC-100k and SEC-F8-F11. (F) Viral titer quantified by qPCR. (G) Viral genomes/nanoparticle ratio. SEC F8-F11 displayed much less viral genomes/nanoparticle. Statistical analysis was performed using the unpaired Student’s t-test (* p < 0.05, **p < 0.01). Results are shown as mean ± SEM for UC-100k (N = 3) and SEC F8-F11 (N = 3).

**Figure 11 - EV-AAVs mediate higher transduction efficiency than solo AAVs, particularly when purified by SEC**

(A) Schematic representation of the transduction assay. Neuro 2A (N2a) cells were transduced at a MOI of 10 000 with EV-AAVs isolated by UC-20k, UC-100k, and SEC (F8-F11). N2a cells transduced with solo AAV1/2 at a MOI of 10 000 were used as control. GFP expression was measured 48h post-infection by FACS analysis (B) and confirmed by ICC (C). EV-AAVs purified by SEC were the most efficient. Statistical analysis relative to control was performed using Ordinary One-way ANOVA with Dunnett’s post-hoc test (*p<0.05, ****p< 0.0001). Results are shown as mean ± SEM for UC-20k (N=3), UC-100k (N=3), F8-F11 (N=3), AAV (N=1). Scale bar = 200µm. See also Supplementary Fig. 5.

**Figure 12 - Intravenously injected SEC EV-AAVs are able to cross the BBB and transduce wild-type mouse brains**

(A) Representative study design. Postnatal day 1 (P1) WT C57BL/6 mice were intravenously injected with 1×10^10^ vg of EV-AAVs encoding GFP and purified by SEC. Mice were sacrificed at P100 and brains were processed. (B) Representative images of GFP IHC in the brains of P100 mice. SEC purified EV-AAVs were able to transduce WT mouse brains, with preferential tropism to the cerebellum (lobule 9 and 10 (L9/10) shown in detail), brainstem and to the choroid plexus. A non-injected mouse was used as staining control. Cerebellum and brainstem scale bar = 250 µm; Lateral Ventricle scale bar = 500 µm. (C) Representative higher magnification images of cerebellum, brainstem, lateral and 4^th^ ventricles from GFP IHC in the brains of P100 mice. Cerebellum, brainstem, lateral and 4^th^ ventricles scale bar = 50 µm.. WT P100 EV-AAVs N = 4.

**Supplementary Figure 7 - EV-enriched fractions present low protein contamination** (A) To confirm that free proteins were not co-eluted with EVs by SEC, western blots were performed. The same volume of each SEC fraction was loaded, and membranes were stained with Ponceau. (B) Protein bands corresponding to “contaminating proteins” were detected between F12 and F26. (C) The protein elution profile was confirmed by BCA. (B, C) Results are shown as mean ± SEM for Ponceau staining (N = 3) and BCA (N = 2).

**Supplementary Figure 8 - Uncropped western blot from main figure** (A) Uncropped western blot from the corresponding cropped western blot shown within the main text.

**Supplementary Figure 9 – Characterization of SEC F7 to F12 by negative Transmission Electron Microscopy (TEM)** (A) SEC fractions (F7 to F12) were imaged by negative stain TEM. Scale bar = 50nm.

**Supplementary Figure 10 - EV-AAVs (F7 to F11) were more efficient at transducing neuronal cells than solo AAVs (F13) (**A) Neuro 2A cells were transduced individually at a MOI of 10 000 with SEC fractions F7 to F13. Non-transduced cells were used as control (CTRL). GFP expression was evaluated 48h post-transduction by ICC. Representative fluorescence images stained with anti-GFP (green) antibody and co-labeled with DAPI (blue). Scale bar = 200 µm.

**Supplementary Figure 11 - Two different populations of EVs are obtained during ultracentrifugation of AAV production medium at 20 000 g (UC-20k) and 100 000 g (UC-100k)** (A) Schematic representation of the extracellular vesicle-associated adeno-associated virus (EV-AAV) isolation protocol by ultracentrifugation (UC). AAV production medium was submitted to two steps of ultracentrifugation (1h at 20k g followed by 1h30 at 100k g). In both steps, pelleted material was collected and subsequently analyzed. (B) The presence of EV protein markers (FLOT-1, AAV capsid proteins (VP1, VP2, VP3) and cell contamination markers (CNX) was analyzed by Western Blot. UC-20k and UC-100k were loaded in the same proportion. FLOT-1 and AAV capsid proteins were present in UC-20k and UC-100k.CNX was mostly present on UC-20k. Relative expression of AAV capsid proteins, FLOT-1 and CNX normalized to 20k fraction was quantified (C). AAV capsid proteins were significantly enriched in UC-100k fraction. AAV purified by FPLC and HEK293T cell lysates were used as controls. (D, E, F) Nanoparticle quantity and size distribution were determined by Nanoparticle Tracking Analysis (NTA). No significant differences were observed regarding nanoparticles concentration. Nanoparticles population was smaller in the UC-100k fraction (UC-20k: 116.3 nm ± 6.093; UC-100k: 80.90 nm ± 2.011). (G) Viral titter was quantified by quantitative polymerase chain reaction (qPCR). (H) Number of viral genomes per nanoparticle showed a tendency to be higher in UC-100k. (C - H) Statistical analysis was performed using the unpaired Student’s t-test (** p < 0.01). Results are shown as mean ± SEM for UC-20k (N = 3), UC-100k (N = 3), except in (G) qPCR (N = 9) per group.

**Supplementary Figure 12 - EV-AAVs are more efficient at transduction of cells than solo AAVs, particularly when purified by SEC (**A) Neuro 2A (N2a) cells were transduced at a MOI of 10 000 with UC-20k, UC-100k, and SEC-F8-F11. N2a cells transduced with solo AAV1/2 were used as control. GFP expression was evaluated 48h post-infection by ICC. Representative fluorescence images stained with anti-GFP (green) antibody and co-labeled with DAPI (blue). Scale bar = 200 µm.

## References

1. Ghosh, S., Brown, A.M., Jenkins, C., and Campbell, K. (2020). Viral Vector Systems for Gene Therapy: A Comprehensive Literature Review of Progress and Biosafety Challenges. Appl. Biosaf. 25, 7–18. 10.1177/1535676019899502.

2. Atchison, R.W., Casto, B.C., and Hammon, W.M.D. (1965). Adenovirus-Associated Defective Virus Particles. Science (80-.). 149, 754–756. 10.1126/SCIENCE.149.3685.754.

3. Saraiva, J., Nobre, R.J., and Pereira de Almeida, L. (2016). Gene therapy for the CNS using AAVs: The impact of systemic delivery by AAV9. J. Control. Release 241, 94–109. 10.1016/j.jconrel.2016.09.011.

4. Ozelo, M.C., Mahlangu, J., Pasi, K.J., Giermasz, A., Leavitt, A.D., Laffan, M., Symington, E., Quon, D. V, Wang, J.-D., Peerlinck, K., et al. (2022). Valoctocogene Roxaparvovec Gene Therapy for Hemophilia A. N. Engl. J. Med. 386, 1013–1025. 10.1056/NEJMoa2113708.

5. Tai, C.-H., Lee, N.-C., Chien, Y.-H., Byrne, B.J., Muramatsu, S.-I., Tseng, S.-H., and Hwu, W.-L. (2022). Long-term efficacy and safety of eladocagene exuparvovec in patients with AADC deficiency. Mol. Ther. 30, 509–518. 10.1016/j.ymthe.2021.11.005.

6. Maguire, A.M., Simonelli, F., Pierce, E.A., Pugh, E.N.J., Mingozzi, F., Bennicelli, J., Banfi, S., Marshall, K.A., Testa, F., Surace, E.M., et al. (2008). Safety and efficacy of gene transfer for Leber’s congenital amaurosis. N. Engl. J. Med. 358, 2240–2248. 10.1056/NEJMoa0802315.

7. Mendell, J.R., Al-Zaidy, S., Shell, R., Arnold, W.D., Rodino-Klapac, L.R., Prior, T.W., Lowes, L., Alfano, L., Berry, K., Church, K., et al. (2017). Single-Dose Gene-Replacement Therapy for Spinal Muscular Atrophy. N. Engl. J. Med. 377, 1713–1722. 10.1056/nejmoa1706198.

8. Heo, Y.-A. (2023). Etranacogene Dezaparvovec: First Approval Etranacogene Dezaparvovec (Hemgenix ®): Key Points. Drugs 83, 347–352. 10.1007/s40265-023-01845-0.

9. Gaudet, D., Méthot, J., Déry, S., Brisson, D., Essiembre, C., Tremblay, G., Tremblay, K., de Wal, J., Twisk, J., van den Bulk, N., et al. (2013). Efficacy and long-term safety of alipogene tiparvovec (AAV1-LPLS447X) gene therapy for lipoprotein lipase deficiency: an open-label trial. Gene Ther. 20, 361–369. 10.1038/gt.2012.43.

10. Colella, P., Ronzitti, G., and Mingozzi, F. (2018). Emerging Issues in AAV-Mediated In Vivo Gene Therapy. Mol. Ther. - Methods Clin. Dev. 8, 87–104. 10.1016/j.omtm.2017.11.007.

11. Louis Jeune, V., Joergensen, J.A., Hajjar, R.J., and Weber, T. (2013). Pre-existing Anti–Adeno-Associated Virus Antibodies as a Challenge in AAV Gene Therapy. Hum. Gene Ther. Methods 24, 59–67. 10.1089/hgtb.2012.243.

12. Huang, L., Wan, J., Wu, Y., Tian, Y., Yao, Y., Yao, S., Ji, X., Wang, S., Su, Z., and Xu, H. (2021). Challenges in adeno-associated virus-based treatment of central nervous system diseases through systemic injection. Life Sci. 270. 10.1016/J.LFS.2021.119142.

13. Maguire, C.A., Balaj, L., Sivaraman, S., Crommentuijn, M.H.W., Ericsson, M., Mincheva-Nilsson, L., Baranov, V., Gianni, D., Tannous, B.A., Sena-Esteves, M., et al. (2012). Microvesicle-associated AAV vector as a novel gene delivery system. Mol. Ther. 20, 960–971. 10.1038/mt.2011.303.

14. György, B., Fitzpatrick, Z., Crommentuijn, M.H.W., Mu, D., and Maguire, C.A. (2014). Naturally enveloped AAV vectors for shielding neutralizing antibodies and robust gene delivery invivo. Biomaterials 35, 7598–7609. 10.1016/j.biomaterials.2014.05.032.

15. Cheng, M., Dietz, L., Gong, Y., Eichler, F., Nammour, J., Ng, C., Grimm, D., and Maguire, C.A. (2021). Neutralizing Antibody Evasion and Transduction with Purified Extracellular Vesicle-Enveloped Adeno-Associated Virus Vectors. Hum. Gene Ther. 32, 1457–1470. 10.1089/hum.2021.122.

16. Meliani, A., Boisgerault, F., Fitzpatrick, Z., Marmier, S., Leborgne, C., Collaud, F., Simon Sola, M., Charles, S., Ronzitti, G., Vignaud, A., et al. (2017). Enhanced liver gene transfer and evasion of preexisting humoral immunity with exosome-enveloped AAV vectors. Blood Adv. 1, 2019– 2031. 10.1182/bloodadvances.2017010181.

17. Wang, W., Liu, J., Yang, M., Qiu, R., Li, Y., Bian, S., Hao, B., and Lei, B. (2021). Intravitreal injection of an exosome-associated adeno-associated viral vector enhances retinoschisin 1 gene transduction in the Mouse Retina. Hum. Gene Ther. 32, 707–716. 10.1089/hum.2020.328.

18. Hudry, E., Martin, C., Gandhi, S., György, B., Scheffer, D.I., Mu, D., Merkel, S.F., Mingozzi, F., Fitzpatrick, Z., Dimant, H., et al. (2016). Exosome-associated AAV vector as a robust and convenient neuroscience tool. Gene Ther. 23, 380–392. 10.1038/gt.2016.11.

19. Liu, B., Li, Z., Huang, S., Yan, B., He, S., Chen, F., and Liang, Y. (2021). AAV-Containing Exosomes as a Novel Vector for Improved Gene Delivery to Lung Cancer Cells. Front. Cell Dev. Biol. 9, 1–12. 10.3389/fcell.2021.707607.

20. Volak, A., LeRoy, S.G., Natasan, J.S., Park, D.J., Cheah, P.S., Maus, A., Fitzpatrick, Z., Hudry, E., Pinkham, K., Gandhi, S., et al. (2018). Virus vector-mediated genetic modification of brain tumor stromal cells after intravenous delivery. J. Neurooncol. 139, 293–305. 10.1007/s11060-018-2889-2.

21. Hong, C.S., Funk, S., Muller, L., Boyiadzis, M., and Whiteside, T.L. (2016). Isolation of biologically active and morphologically intact exosomes from plasma of patients with cancer. J. Extracell. Vesicles 5. 10.3402/jev.v5.29289.

22. Lobb, R.J., Becker, M., Wen, S.W., Wong, C.S.F., Wiegmans, A.P., Leimgruber, A., and Möller, A. (2015). Optimized exosome isolation protocol for cell culture supernatant and human plasma. J. Extracell. Vesicles 4. 10.3402/jev.v4.27031.

23. Soares, M., Pinto, M.M., Nobre, R.J., de Almeida, L.P., da Graça Rasteiro, M., Almeida-Santos, T., Ramalho-Santos, J., and Sousa, A.P. (2023). Isolation of Extracellular Vesicles from Human Follicular Fluid: Size-Exclusion Chromatography versus Ultracentrifugation. Biomolecules 13. 10.3390/biom13020278.

24. Gaspar, L.S., Santana, M.M., Henriques, C., Pinto, M.M., Ribeiro-Rodrigues, T.M., Girão, H., Nobre, R.J., and Pereira de Almeida, L. (2020). Simple and Fast SEC-Based Protocol to Isolate Human Plasma-Derived Extracellular Vesicles for Transcriptional Research. Mol. Ther. - Methods Clin. Dev. 18, 723–737. 10.1016/j.omtm.2020.07.012.

25. Chen, W., Yao, S., Wan, J., Tian, Y., Huang, L., Wang, S., Akter, F., Wu, Y., Yao, Y., and Zhang, X. (2021). BBB-crossing adeno-associated virus vector: An excellent gene delivery tool for CNS disease treatment. J. Control. Release 333, 129–138. 10.1016/j.jconrel.2021.03.029.

26. Grieger, J.C., Choi, V.W., and Samulski, R.J. (2006). Production and characterization of adeno-associated viral vectors. Nat. Protoc. 1, 1412–1428. 10.1038/nprot.2006.207.

27. Vergauwen, G., Dhondt, B., Van Deun, J., De Smedt, E., Berx, G., Timmerman, E., Gevaert, K., Miinalainen, I., Cocquyt, V., Braems, G., et al. (2017). Confounding factors of ultrafiltration and protein analysis in extracellular vesicle research. Sci. Rep. 7, 1–12. 10.1038/s41598-017-02599-y.

28. György, B., Sage, C., Indzhykulian, A.A., Scheffer, D.I., Brisson, A.R., Tan, S., Wu, X., Volak, A., Mu, D., Tamvakologos, P.I., et al. (2017). Rescue of Hearing by Gene Delivery to Inner-Ear Hair Cells Using Exosome-Associated AAV. Mol. Ther. 25, 379–391. 10.1016/j.ymthe.2016.12.010.

29. Wu, N., and Ataai, M.M. (2000). Production of viral vectors for gene therapy applications. Curr. Opin. Biotechnol. 11, 205–208. 10.1016/S0958-1669(00)00080-X.

30. Nobre, R.J., Almeida, L.P. de, Jorge Nobre, R., and Pereira de Almeida, L. (2011). Gene Therapy for Parkinsons and Alzheimers Diseases: from the Bench to Clinical Trials. Curr. Pharm. Des. 17, 3434–3445. 10.2174/138161211798072472.

31. Choudhury, S.R., Hudry, E., Maguire, C.A., Sena-Esteves, M., Breakefield, X.O., and Grandi, P. (2017). Viral vectors for therapy of neurologic diseases. Neuropharmacology 120, 63–80. 10.1016/j.neuropharm.2016.02.013.

32. Rufino-Ramos, D., Albuquerque, P.R., Carmona, V., Perfeito, R., Nobre, R.J., and Pereira de Almeida, L. (2017). Extracellular vesicles: Novel promising delivery systems for therapy of brain diseases. J. Control. Release 262, 247–258. 10.1016/j.jconrel.2017.07.001.

33. Ratajczak, J., Wysoczynski, M., Hayek, F., Janowska-Wieczorek, A., and Ratajczak, M.Z. (2006). Membrane-derived microvesicles: Important and underappreciated mediators of cell-to-cell communication. Leukemia 20, 1487–1495. 10.1038/sj.leu.2404296.

34. Frühbeis, C., Fröhlich, D., Kuo, W.P., Amphornrat, J., Thilemann, S., Saab, A.S., Kirchhoff, F., Möbius, W., Goebbels, S., Nave, K.A., et al. (2013). Neurotransmitter-Triggered Transfer of Exosomes Mediates Oligodendrocyte-Neuron Communication. PLoS Biol. 11. 10.1371/journal.pbio.1001604.

35. Alvarez-Erviti, L., Seow, Y., Yin, H., Betts, C., Lakhal, S., and Wood, M.J.A.A. (2011). Delivery of siRNA to the mouse brain by systemic injection of targeted exosomes. Nat. Biotechnol. 29, 341–345. 10.1038/nbt.1807.

36. Wassmer, S.J., Carvalho, L.S., György, B., Vandenberghe, L.H., and Maguire, C.A. (2017). Exosome-associated AAV2 vector mediates robust gene delivery into the murine retina upon intravitreal injection. Sci. Rep. 7, 45329. 10.1038/srep45329.

37. Khan, N., Maurya, S., Bammidi, S., and Jayandharan, G.R. (2020). AAV6 Vexosomes Mediate Robust Suicide Gene Delivery in a Murine Model of Hepatocellular Carcinoma. Mol. Ther. - Methods Clin. Dev. 17, 497–504. 10.1016/j.omtm.2020.03.006.

38. Orefice, N.S., Souchet, B., Braudeau, J., Alves, S., Piguet, F., Collaud, F., Ronzitti, G., Tada, S., Hantraye, P., Mingozzi, F., et al. (2019). Real-Time Monitoring of Exosome Enveloped-AAV Spreading by Endomicroscopy Approach: A New Tool for Gene Delivery in the Brain. Mol. Ther. - Methods Clin. Dev. 14, 237–251. 10.1016/j.omtm.2019.06.005.

39. Maurya, S., and Jayandharan, G.R. (2020). Exosome-associated SUMOylation mutant AAV demonstrates improved ocular gene transfer efficiency in vivo. Virus Res. 283. 10.1016/j.virusres.2020.197966.

40. Benedikter, B.J., Bouwman, F.G., Vajen, T., Heinzmann, A.C.A., Grauls, G., Mariman, E.C., Wouters, E.F.M., Savelkoul, P.H., Lopez-Iglesias, C., Koenen, R.R., et al. (2017). Ultrafiltration combined with size exclusion chromatography efficiently isolates extracellular vesicles from cell culture media for compositional and functional studies. Sci. Rep. 7, 15297. 10.1038/s41598-017-15717-7.

41. Maruno, T., Usami, K., Ishii, K., Torisu, T., and Uchiyama, S. (2021). Comprehensive Size Distribution and Composition Analysis of Adeno-Associated Virus Vector by Multiwavelength Sedimentation Velocity Analytical Ultracentrifugation. J. Pharm. Sci. 110, 3375–3384. 10.1016/J.XPHS.2021.06.031.

42. Rufino-Ramos, D., Lule, S., Mahjoum, S., Ughetto, S., Cristopher Bragg, D., Pereira de Almeida, L., Breakefield, X.O., and Breyne, K. (2022). Using genetically modified extracellular vesicles as a non-invasive strategy to evaluate brain-specific cargo. Biomaterials 281, 121366. 10.1016/J.BIOMATERIALS.2022.121366.

43. Webber, J., and Clayton, A. (2013). How pure are your vesicles? J. Extracell. Vesicles 2, 1–6. 10.3402/jev.v2i0.19861.

44. Kumar, P., Wu, H., McBride, J.L., Jung, K.-E., Hee Kim, M., Davidson, B.L., Kyung Lee, S., Shankar, P., and Manjunath, N. (2007). Transvascular delivery of small interfering RNA to the central nervous system. Nature 448, 39–43. 10.1038/nature05901.

45. Rufino-Ramos, D., Albuquerque, P.R., Leandro, K., Carmona, V., Martins, I.M., Fernandes, R., Henriques, C., Lobo, D., Faro, R., Perfeito, R., et al. (2023). Extracellular vesicle-based delivery of silencing sequences for the treatment of Machado-Joseph disease/spinocerebellar ataxia type 3. Mol. Ther. 31, 1275–1292. 10.1016/j.ymthe.2023.04.001.

46. Yamazaki, Y., Hirai, Y., Miyake, K., and Shimada, T. (2014). Targeted gene transfer into ependymal cells through intraventricular injection of AAV1 vector and long-term enzyme replacement via the CSF. Sci. Rep. 4, 1–7. 10.1038/srep05506.

47. Conceição, M., Costa, P., Conceiç, M., Hirai, H., Pereira, L., Almeida, D., Mendonça, L., Nóbrega, C., Gomes, C., Costa, P., et al. (2016). Intravenous administration of brain-targeted stable nucleic acid lipid particles alleviates Machado-Joseph disease neurological phenotype. Biomaterials 82, 124–137. 10.1016/j.biomaterials.2015.12.021.

48. Paulson, H.L., Shakkottai, V.G., Clark, H.B., and Orr, H.T. (2017). Polyglutamine spinocerebellar ataxias-from genes to potential treatments. Nat. Rev. Neurosci. 18, 613–626. 10.1038/nrn.2017.92.

49. Lopes, M.M., Paysan, J., Rino, J., Lopes, S.M., Pereira de Almeida, L., Cortes, L., and Nobre, R.J. (2022). A new protocol for whole-brain biodistribution analysis of AAVs by tissue clearing, light-sheet microscopy and semi-automated spatial quantification. Gene Ther. 2022 2912 29, 665–679. 10.1038/s41434-022-00372-z.

50. Wang, D., Li, S., Gessler, D.J., Xie, J., Zhong, L., Li, J., Tran, K., Van Vliet, K., Ren, L., Su, Q., et al. (2018). A Rationally Engineered Capsid Variant of AAV9 for Systemic CNS-Directed and Peripheral Tissue-Detargeted Gene Delivery in Neonates. Mol. Ther. - Methods Clin. Dev. 9, 234–246. 10.1016/j.omtm.2018.03.004.

51. Byrne, L.C., Lin, Y.J., Lee, T., Schaffer, D. V., and Flannery, J.G. (2015). The expression pattern of systemically injected AAV9 in the developing mouse retina is determined by age. Mol. Ther. 23, 290–296. 10.1038/mt.2014.181.

52. Huda, F., Konno, A., Matsuzaki, Y., Goenawan, H., Miyake, K., Shimada, T., and Hirai, H. (2014). Distinct transduction profiles in the CNS via three injection routes of AAV9 and the application to generation of a neurodegenerative mouse model. Mol. Ther. - Methods Clin. Dev. 1, 14032. 10.1038/mtm.2014.32.

53. Lamichhane, T.N., Raiker, R.S., and Jay, S.M. (2015). Exogenous DNA Loading into Extracellular Vesicles via Electroporation is Size-Dependent and Enables Limited Gene Delivery. Mol. Pharm. 12, 3650–3657. 10.1021/acs.molpharmaceut.5b00364.

54. Thierry, Théry, C., Amigorena, S., Raposo, G., and Clayton, A. (2006). Isolation and Characterization of Exosomes from Cell Culture Supernatants. Curr. Protoc. Cell Biol. Chapter 3, 1–29. 10.1002/0471143030.cb0322s30.

